# Retinoic acid signaling guides the efficiency of inner ear organoid-genesis and governs sensory-nonsensory fate specification

**DOI:** 10.1101/2025.03.21.644434

**Authors:** R. Keith Duncan, Liqian Liu, Mo Moyer, Andrew Wylie, Ranya Dano, Luis Cassinotti

## Abstract

Inner ear organoid development—from germ layer to otocyst formation—relies on timed chemical cues to recapitulate major signals *in vivo.* In contrast, later stages of differentiation—from otic vesicle (OV) to organoid formation—are self-guided, even though these stages are modulated by several key morphogens *in vivo*. We sought to elucidate additional morphogens that might improve culture efficiency and influence cell fate decisions. Using a whole-transcriptomic approach, we identified major differences in native and stem cell-derived OVs related to anterior-posterior patterning and retinoic acid (RA) signaling. Increasing the level of RA during OV formation in these cultures modulated organoid efficiency, increased nonsensory markers, decreased sensory markers, and decreased hair cell production. The organoid culture platform mimics the exquisite RA sensitivity found in normal inner ear development and may help identify RA-responsive genes driving organogenesis and cell fate specification.

## INTRODUCTION

Three-dimensional organoid culture systems provide a unique parallel approach to *in vivo* inner ear research. These culture systems are accessible throughout the differentiation process, scalable to high-throughput technologies, and easy to manipulate, opening new paths to study development, disease, therapeutics, and regeneration. To reach their full potential, inner ear organoids (IEOs) must mimic the complexity of inner ear organs and become malleable enough to control cell fate decisions. Recent advances have underscored the capabilities of mouse- and human-derived IEOs to follow the developmental trajectories of native tissue^1–3^, explored mechanisms of disease^4,5^, and continued to refine and optimize benchmark protocols^6^. A fundamental principle of these protocols is that a basic understanding of signaling mechanisms active in early embryonic development should serve as the foundation for guiding each step in differentiation.

To date, the vast majority of IEO culture paradigms include a cue-driven phase followed by self- assembly into complex structures. In the cue-drive phase, exogenous chemicals are applied to sequentially guide a three-dimensional spheroid of pluripotent stem cells toward the formation of ectodermal otic intermediates^7,8^. Following the initial aggregation into spheres, an inhibitor of the TGFβ pathway directs differentiation to favor ectoderm and limit endoderm and mesoderm^9,10^. Additional drug cues further refine the lineage path toward non-neural ectoderm, preplacodal domains, otic placode, and ultimately formation of otic vesicles (OVs) within the larger spheroid. Almost all sensory and nonsensory cells within the six distinct inner ear organs are derived from the multipotent epithelial progenitor cells of the OV^11^. In IEO culture paradigms, the formation of OVs within the spheroid typically marks the transition to a self-assembly phase without the addition of exogenous morphogens to further guide development.

*In vivo*, the axial specification of the inner ear and the corresponding spatial organization of the statoacoustic organs are largely defined during morphogenesis of the OV to a mature inner ear, a developmental time period that matches the self-assembly phase of IEOs. Diffusible signals from neighboring tissues—the hindbrain, periotic mesenchyme, notochord, and floor plate—sequentially pattern the inner ear along anterior-posterior, dorsal-ventral, and medial-lateral axes^11,12^. Axial specification is progressive, beginning with an anterior-posterior asymmetry that establishes the neural-sensory domain in the anterior region and nonsensory tissues in the posterior region, save for the posterior crista^12^. Following anterior-posterior segmentation, opposing gradients of WNT and SHH establish dorsal-ventral domains that largely define the separation of vestibular-auditory organs, respectively. Medial-lateral patterning appears important for neural segmentation and shifting sensory domains during maturation, but the underlying mechanisms are unclear^12–14^.

While recent reports have explored cochlear-vestibular patterning in organoids by manipulating WNT and SHH signaling^6^, there has been no attempt to systematically examine the early stage of anterior-posterior patterning through the actions of retinoic acid (RA). RA signaling is indispensable in inner ear development. As the otocyst forms *in vivo*, an anterior-to-posterior wave of RA activity defines sensory-nonsensory compartments^15^. During a narrow critical window of otocyst development, low RA exposure in anterior domains guides a prosensory fate, and higher exposure in posterior domains induces nonsensory fates. RA effects are pleiotropic and involve a large number of RA-responsive genes^16^, but only a few definitive RA-target genes have been identified in ear development^15,17^. Notably, RA dysregulation *in vivo*, whether from genetics or environmental factors, is teratogenic; both excess and deficiency cause ear abnormalities and even complete arrest of inner ear development^17^. We hypothesize that RA plays equally important roles in organoid-genesis and in controlling sensory (anterior) versus nonsensory (posterior) fates *in vitro*.

In this study, we fine tune the IEO protocol with RA to systematically modulate organoid production and develop RA supplementation as a tool for shifting between sensory to nonsensory cell fates. An examination of the transcriptome of isolated OVs from embryonic mouse compared to that of age-matched organoid cultures confirmed dysregulation of anterior-posterior pattern specification in the cultures and pointed to the involvement of RA in contributing to organoid heterogeneity and dysmorphogenesis, where some OVs fail to mature into large cystic organoids. Our data replicate the exquisite sensitivity of early stages of inner ear development to spatiotemporal perturbations in RA level, and definitively show that systematic variations in RA level tip the balance from sensory to nonsensory domain formation. Consequently, gaining control over RA signaling may lead to rationale approaches for enhancing hair cell production or producing specific populations of nonsensory cell types. These observations establish RA as another key morphogen in IEO paradigms and pave the way toward a model of axial patterning in future organoid research. Understanding the effects of RA signaling and optimizing RA supplementation during formation of IEOs will be crucial to produce the large quantities of hair cells required for future downstream applications.

## RESULTS

### R1/E mouse ESCs produce otic vesicle-like and organoid structures

To produce derived OVs and IEOs, we used a protocol optimized for R1/E mouse embryonic stem cells (ESCs) published previously^18^ (**Figure 1A**), referring to this culture paradigm as BSFL for the compounds added in the cue-driven phase (i.e., <u>B</u>MP4/<u>S</u>B431542 treatment at day 3 (D3) and <u>F</u>GF2/<u>L</u>DN193189 treatment at D4.5). These cultures recapitulated major checkpoints for IEO production, from formation of the initial spheroid (**Figure 1B**) to production of a ruffling non-neural ectodermal layer **(Figure 1C**), and generation of Pax2-positive otic vesicle-like structures by D12 (**Figure 1D**). These OVs could be released from the surrounding aggregate by a combination of enzymatic and mechanical disruption (**Figure 1E**), facilitating downstream analysis of their transcriptomic profiles. Within intact aggregates, the vesicles themselves expanded in size by D20, often becoming large, protruding fluid-filled cysts (**Figure 1F**). Immunofluorescence staining of epithelia dissected from these cysts revealed MyoVIIa-positive hair cell-like cells, indicating sensory domains within the IEO (**Figure 1G**).

**Figure 1.**
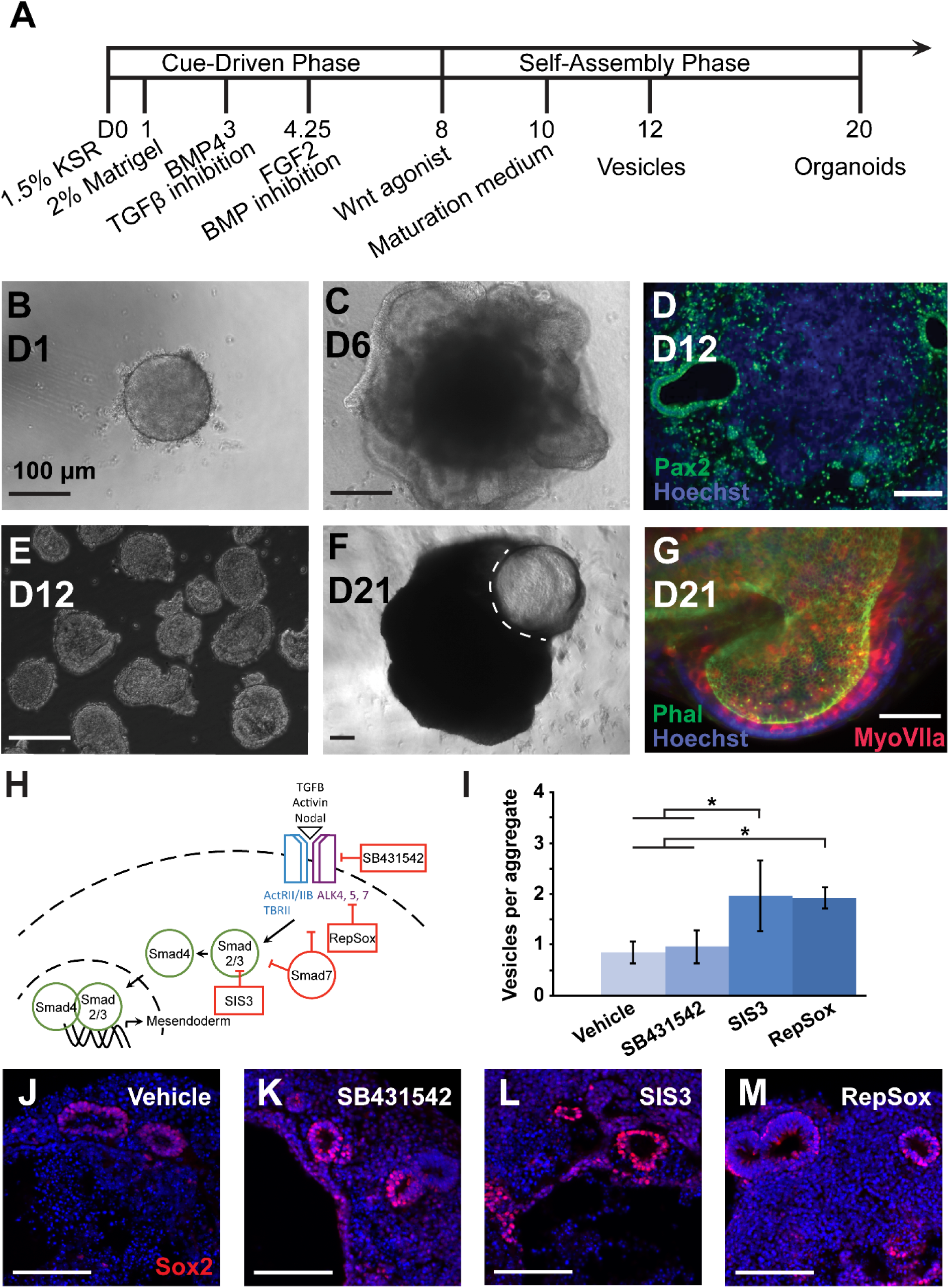
Generation of inner ear organoids. (A) The timeline of IEO formation is broadly broken into a cue-driven phase sequentially driving generation of definitive ectoderm with the addition of extracellular matrix (Matrigel), non-neural ectoderm with BMP4 and TGFβ inhibition, and otic placode with a combination of FGF2 and BMP inhibition followed by WNT activation. Late-stage development is marked by a self-assembly phase producing otic veiscles and ultimately IEOs. (B) Spheroids form in ultra-low attachment wells by D1 followed by (C) extensive ruffling of an outer epithelial later of non-neural ectoderm by D6. (D) Thin sections through D12 aggregates reveal internal, Pax2-positive OVs (green; nuclear counterstain Hoechst in blue). (E) The OVs can be isolated by a combination of enzymatic digestion and mechanical trituration. (F) Protruding (dotted outline) organoid cysts can be identified after D20. (G) Microdissected organoids contain patches of sensory epithelia, where phalloiodin-labeled hair bundles (green) extended into the lumen from MyoVIIa-positive hair cells (red; nuclear counterstain Hoechst in blue). (H) A schematic shows mechanisms of TGFβ pathway inhibition. (I) The efficiency of OV production was quantified at D12 in aggregates treated with different TGFβ inhibitors on culture D3 (* *P*<0.05; mean ± one standard deviation). N = 8 (vehicle), 8 (SB431542), 5 (SIS3), 5 (RepSox). (J-M) Example immunofluorescence images of cyrosections from D12 aggregates in each condition stained with the OV marker Sox2 (red) and counterstained with Hoechst (blue). Scale bars: 100 μm.

### TGF**β** inhibition is dispensable for OV production but indispensable for generating IEOs

The ability to fine-tune organoid efficiency and fate requires additional insights into the molecular mechanisms shaping organoid-genesis. Efficiency is linked to each major developmental milestone in the procedure. For example, organoid production is tightly controlled by the dose and timing of exogenous BMP and TGFβ inhibition in order to generate definitive ectoderm while suppressing mesoendodermal fates^2,8^. In devising a scheme to compare the whole transcriptomes of derived and native OVs, we first sought to identify conditions that resulted in similar degrees of OV production but varied degrees of organoid-genesis, focusing our attention on TGFβ inhibition at D3. Otic vesicles were collected from cultures treated on D3 with vehicle control media and one of several TGFβ inhibitors (i.e. SB431542, SIS3, and RepSox). Whereas SB431542 and RepSox block signaling at the receptor level upstream of effectors Smad2 and Smad3, SIS3 specifically blocks activation of Smad3 (**Figure 1H**). SB431542-treated and untreated aggregates produced vesicles at comparable efficiency; however, RepSox and SIS3 resulted in significantly more vesicles per aggregate (*P*<0.05) (**Figure 1I**). Despite differential efficiency in vesicle formation, the vesicles in all conditions were qualitatively similar in size and expression of native OV marker Sox2 (**Figure 1J-M**). Vesicles in all samples were also positive for otic markers Six1, Ecad, and Pax2 (data not shown). Organoids form in the late maturation phase through the continued differentiation of vesicles; however, not all vesicles will necessarily become organoids. Large, fluid-filled organoid cysts were observed in the SB431542 (11/12 trials), RepSox (7/7 trials), and SIS3 treatment groups (6/7 trials). No statistically reliable difference was observed amongst the 3 inhibitor groups when quantifying the percentage of aggregates with one or more visible IEO (**see Figure S1A**), but for those aggregates capable of producing organoid cysts, RepSox appeared to produce more per aggregate than SB431542 though this was not quantified. Notably, the vehicle control group never produced internal or protruding cysts (0/8 trials). Immunofluorescence verified hair cell production in all inhibitor conditions (**see Figure S1B-D**), but hair cells were never observed in cultures without TGFβ inhibition.

### Differences in the transcriptomes of derived compared to native OVs involve anterior-posterior fate specification

Based on these observations, we chose to compare the transcriptomes of derived OVs cultured in BSFL conditions with those in BFL conditions, where the TGFβ inhibitor was omitted. Both were benchmarked against E10.5 OVs harvested from C57BL/6 embryos. The D12 and E10.5 time points were considered analogous, since in each condition vesicles have just formed and neuroblasts are in the process of delaminating^19,20^. TruSeq cDNA libraries were generated for 3 to 4 independent samples of pooled OVs from each experimental group and examined by RNASeq on an Illumina HiSeq-2500 platform. During quality control steps in the bulk RNASeq pipeline, over 90% of base calls passed filtering (i.e., Illumina onboard chastity filter) and had quality scores of Q30 or greater (**see Table S1**). The alignment of sequences to Ensembl annotated transcripts was performed using HTSeq Count, where at least 96.5% of reads per sample mapped to a feature and at least 86.6% mapped to a single feature (**see Table S2**).

In total, over 22,000 genes had measurable expression levels in these samples, and over 10,000 differentially expressed genes (DEGs) were identified in pairwise comparisons between E10.5, BFL, and BSFL groups (**see Table S3**). To assess variability between individual samples, principal component analysis was performed using the DEGs as features (**Figure 2A**). Overall, the three groups organized into distinct clusters by experimental condition, and 89% of the variance in these samples was captured by the first two components. Clusters formed by BFL and BSFL samples showed a greater spread compared to E10.5 otocysts, reflecting greater inter-experimental variability in the cell cultures as each sample was composed of a pool of 30 to 50 vesicles from single culture plates. A heatmap from hierarchical clustering illustrated the close relationship between BSFL and BFL samples whereas E10.5 OV samples were more distant (**see Figure S2A**). A Venn diagram was generated to summarize the overlap between lists of DEGs generated from pairwise comparisons (**Figure 2B**). Differential expression was determined at fold-change expression above 1.5 (i.e., Log_2_(FC)>0.6), and significance was determined at a *P*-value adjusted for false discovery rate below 0.05 (*AdjP*<0.05). The number of genes unique to the E10.5-BFL comparison was the highest at 1,977, with 1,244 unique to E10.5-BSFL and 256 unique to BSFL-BFL. Additional sets of genes appeared in the union of 2 or 3 comparisons; unsurprisingly, a large number of DEGs (over 4,000) were common to the E10.5-BFL and E10.5-BSFL comparison, reflecting a fundamental difference between *in vivo* and *in vitro* samples.

**Figure 2.**
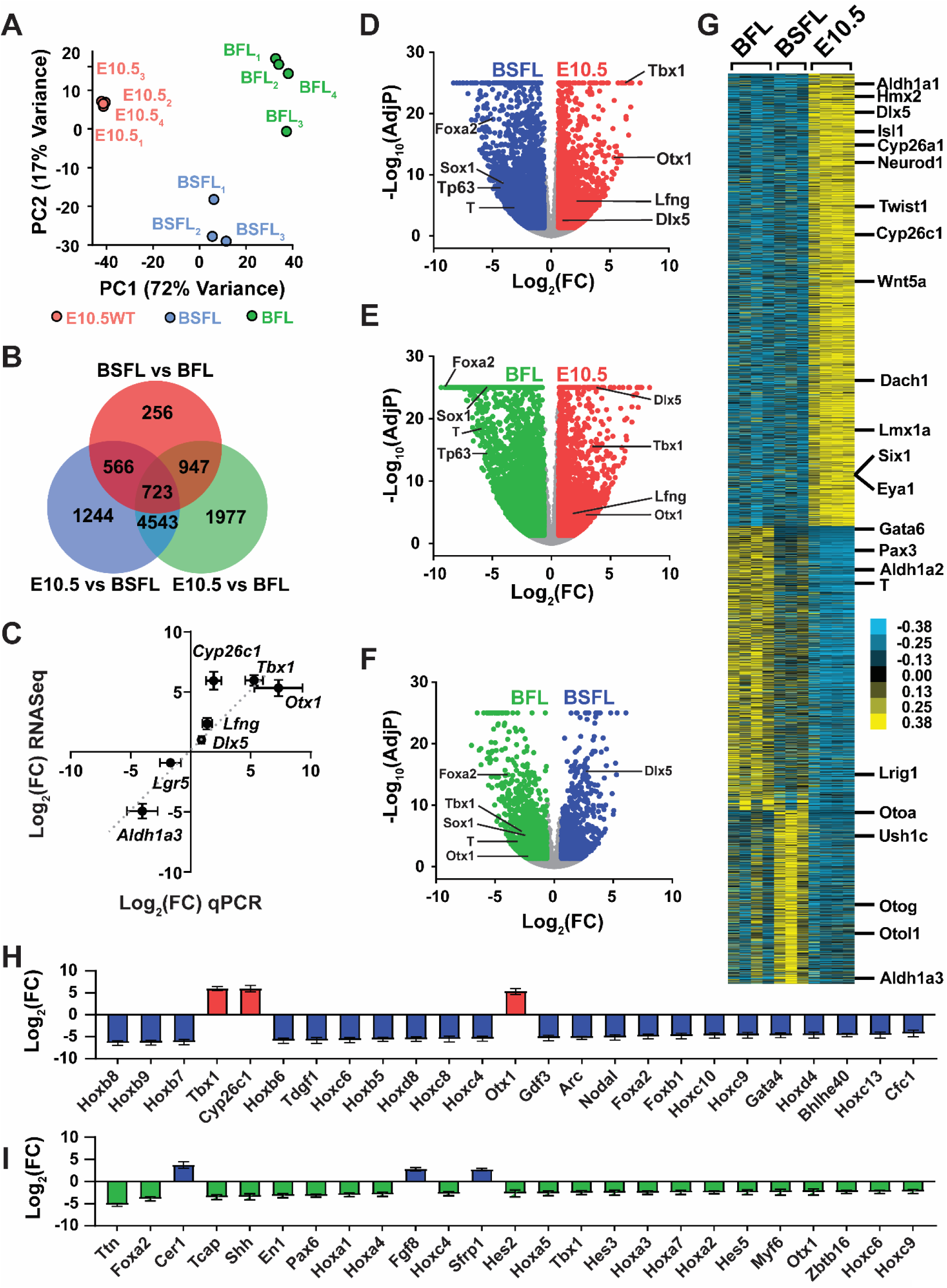
Bulk RNASeq analysis of native E10.5 OVs compared with stem cell-derived OVs at D12 in BSFL and BFL cultures. (A) Principal component analysis illustrates clustering of samples by condition, with some greater separation within culture conditions. (B) Venn diagram shows the DEGs identified in pairwise comparisons of all conditions, including those DEGs unique to each comparison and those at intersections between these comparisons. Over 10,000 DEGs were identified with nearly half of these being common to the pairwise comparison of E10.5 vesicles with either culture condition. (C) The fold-change (Log_2_(FC)) for select genes in RNASeq data is compared to fold-change assessed by qPCR for E10.5 otic vesicles relative to BSFL cultures. N = 6 BSFL for qPCR, 3 BSFL for RNASeq, 3 E10.5 for qPCR, 3 E10.5 for RNASeq. Dotted line indicates the line of unity to highlight agreement between the two methodologies. Linear regression fit to the data gave a slope of 0.94 with Pearson correlation of 0.89 and *P*<0.01. (D-F) Volcano plots of DEGs for pairwise comparisons show the distribution of up- and down-regulated genes. A selection of otic placodal (*Lfng*, *Dlx5*, *Otx1*, *Tbx1*), mesodermal-endodermal (*T*, *Foxa2*), neuroectodermal (*Sox1*), and non-neural ectodermal (*Trp63*) genes are indicated in each panel. (G) A heatmap from hierarchical clustering is illustrated with Java TreeView and the position of select otic and mesodermal-endodermal DEGs are indicated. The top 25 DEGs in the “Anterior/posterior pattern specification” GO term for biological processes are illustrated by log fold-change for (H) E10.5 compared to BSFL cultures and (I) BSFL compared to BFL cultures (mean ± one standard deviation).

Differences in gene expression in the E10.5-BSFL condition were validated by qPCR (**Figure 2C**). Several genes involved in otocyst formation were selected for validation. Log_2_(fold change) in qPCR reactions on additional sets of samples was plotted against Log_2_(fold change) from the RNASeq data. A linear regression through these data gave a slope of 0.94 and correlation coefficient (R^2^) of 0.79, which was statistically significant (Pearson’s *P*<0.01).

To categorize the DEGs further, we generated volcano plots of up- and down-regulated genes (**Figure 2D-F and see Figure S2B-C**). Of the 7,076 DEGs in the E10.5-BSFL comparison, 2,746 were more highly expressed in E10.5 otocysts and 4,330 were more highly expressed in the BSFL-derived vesicles (**Figure 2D**). A selection of genes are annotated on the volcano plots representing markers for otic placode (*Lfng*, *Dlx5*, *Otx1*, *Tbx1*), mesodermal-endodermal fate (*T*, *Foxa2*), neuroectoderm (*Sox1*), and non-neural ectoderm/epidermis (*Trp63*)^21–27^. For the E10.5-BSFL and E10.5-BFL comparisons, the otic markers are enriched in the native OVs whereas the mesodermal-endodermal markers and other ectodermal markers are enriched in the derived OVs (**Figure 2D-E**), suggesting incomplete conversion to an otic fate. Consistent with the lack of TGFβ inhibition in BFL samples, the non-otic lineage genes were more highly expressed in BFL than BSFL conditions (**Figure 2F**).

Hierarchical clustering of the DEGs revealed three distinct clusters (**Figure 2G**). In the first (top) cluster, DEGs were more highly expressed in native E10.5 otocysts compared to either culture condition and included biomarkers of the dorsal (e.g., *Dlx5*, *Hmx2*, *Aldh1a1*)^13,28,29^, medial (e.g., *Dach1* and *Eya1*)^30^, and ventromedial (e.g., *Six1*)^31^ otocyst as well as delaminating neuroblasts (e.g., *Isl1* and *Neurod1*)^32^ and periotic mesenchyme (e.g., *Wnt5a*, *Twist1*, *Cyp26c1*)^28,33^. The second largest cluster had DEGs more highly expressed in BFL cultures and was enriched with mesodermal-endodermal markers (e.g., *Gata6*, *Pax3*, *T/Brachyury*)^22,34,35^, a consequence of the lack of TGFβ inhibition. Finally, the third cluster, with DEGs more highly expressed in BSFL, included several genes associated with nonsensory portions of the inner ear, such as glycoproteins that anchor sensory cells to surrounding structures (e.g., *Otoa*, *Otog*, *Otol1*)^36^.

Gene ontology (GO) analysis associated with DEGs in each pairwise comparison returned generic parent terms related to organismal and developmental processes. Filtering using *elim pruning* to minimize dependency on genes in parent GO terms produced a more refined set, and the top 10 are listed for each pairwise comparison in **Table 1**. Several GO terms in the E10.5-BSFL and E10.5-BFL comparisons are related to high levels of transcriptional activity and cell proliferation, with genes such as *Fos* and *Egr1-4*—along with some genes associated with pluripotency (*Pou5f1* and *Nanog*)—highly enriched in the cultured OVs. Other significant GO terms in these comparisons included terms associated with nonneural ectoderm (“keratinocyte differentiation”) and mesoderm (“sarcomere organization”, “skeletal muscle cell differentiation”, “cardiac muscle”), suggesting incomplete programming of otic placodal fates in the derived OVs. However, “anterior/posterior pattern specification” was the top GO term in E10.5-BSFL and E10.5-BFL comparisons. We plotted the top 25 DEGs in this GO term for E10.5-BSFL (**Figure 2H**). Several Hox genes were enriched in BSFL. Hox genes are major effectors of anterior-posterior body patterning, where the expression of four homologous families (Hoxa, Hoxb, Hoxc, Hoxd) are arranged along the neural tube such that lower numbered isoforms (e.g. *Hoxa1*) are expressed in more anterior domains while higher numbers are expressed posteriorly^37^. For the E10.5-BSFL comparison, all Hox DEGs except for Hoxa2 were more highly expressed in derived OVs than native tissue (**see Figure S2B**), suggesting posteriorization in the cultures. Notably, “anterior/posterior pattern specification” also appeared as a top GO term in the BSFL-BFL comparison. This too was largely driven by enrichment of more posterior Hox genes in BFL compared with BSFL, where nearly a third of the DEGs in this term (25/78) were part of the Hox family (**Figure 2I**). In general, normalized counts for this gene family were BFL > BSFL > E10.5.

**Table 1.** Biological Processes Go Terms, “weight pruning” in iPathwayGuide.

| Comparison | GO term | Genes<br>(DEG/ALL) | Adjusted P-value |
| --- | --- | --- | --- |
| E10.5 v BSFL | Cell proliferation | 731/1832 | 1.4E-8 |
|  | Anterior/posterior pattern specification | 116/225 | 1.7E-7 |
|  | Positive regulation of myeloid cell differentiation | 51/88 | 7.2E-7 |
|  | Positive regulation of transcription by RNA polymerase II | 453/1162 | 1.6E-6 |
|  | Embryonic organ morphogenesis | 146/311 | 1.9E-6 |
|  | Cardiac muscle contraction | 63/120 | 7.9E-6 |
|  | Male meiotic nuclear division | 31/49 | 9.3E-6 |
|  | Synapsis | 29/48 | 1.1E-5 |
|  | Keratinocyte differentiation | 52/97 | 1.3E-5 |
|  | EKR1 and EKR2 cascade | 131/296 | 1.3E-5 |
| E10.5 v BFL | Embryonic organ morphogenesis | 179/311 | 6.7E-14 |
|  | Anterior/posterior pattern specification | 169/225 | 5.6E-13 |
|  | Morphogenesis of a branching structure | 140/235 | 7.1E-11 |
|  | Positive regulation of cell proliferation | 426/885 | 1.9E-10 |
|  | Ear development | 132/231 | 3.9E-10 |
|  | Embryonic skeletal system development | 83/134 | 2.3E-9 |
|  | Muscle organ development | 208/405 | 9.3E-9 |
|  | Positive regulation of cell differentiation | 484/1044 | 2.1E-8 |
|  | Positive regulation of transcription by RNA polymerase II | 520/1162 | 3.1E-8 |
|  | Inflammatory response | 294/580 | 3.5E-7 |
| BSFL v BFL | Embryonic organ morphogenesis | 109/307 | 5.0E-23 |
|  | Embryonic skeletal system development | 58/134 | 5.0E-18 |
|  | Anterior/Posterior pattern specification | 78/223 | 3.7E-17 |
|  | Cell morphogenesis involved in differentiation | 187/768 | 1.3E-16 |
|  | Sarcomere organization | 30/51 | 1.8E-14 |
|  | Muscle organ development | 121/401 | 1.1E-13 |
|  | Skeletal system morphogenesis | 79/248 | 1.4E-13 |
|  | Neuron projection guidance | 75/235 | 6.4E-13 |
|  | Adult behavior | 57/172 | 8.3E-12 |
|  | Extracellular matrix organization | 76/249 | 1.5E-11 |

Anterior-posterior patterning involves gradients of RA for segmentation along the body^38,39^ and in the developing otocyst^15^. Significant GO terms related to RA appeared in all group comparisons (marked with * in **Table 1**). Shifting boundaries of RA synthesis and metabolizing enzymes contribute to gradations in Hox gene expression, and we found the largest fold-change in RA-related genes to be the metabolizing enzyme *Cyp26c1* highest in E10.5 and the synthesizing enzyme *Aldh1a3* highest in BSFL (**see Figure S2C**). The lack of Cyp26 and excessive Aldh1a in BSFL cultures suggests higher levels of RA signaling *in vitro* compared to *in vivo* conditions. Hence, fine-tuning RA signaling during the early stages of organoid-genesis could improve efficiency and allow us to tip the balance between anterior-posterior patterns (i.e., sensory-nonsensory fates).

### Excessive and deficient RA signaling reduces the efficiency of organoid-genesis during a critical time window

*In vivo*, exposure to excess RA and inhibition of RA signaling results in OV dysmorphogenesis and often arrest of inner ear development^17^. To determine whether organoid cultures are similarly sensitive to RA signaling, we exposed cultures of R1E mouse ESCs to exogenous all-trans retinoic acid (atRA) or various inhibitors to RA receptors or RA synthesis during media exchanges between culture D8 and D12. For these cultures and those in the remaining studies, we transitioned to using RepSox as the TGFβ inhibitor on D3, since this compound produced more OVs and increased the reliability of organoid production (so-called BRFL cultures). In control conditions, the efficiency of producing OVs and protruding organoids was about 75% and 30%, respectively (**Figure 3A**). Both inhibition of RA receptors (RARs) with the pan-RAR antagonist AGN193109 and the addition of atRA significantly inhibited organoid production (**Figure 3B**). Similarly, inhibition of endogenous RA biosynthesis with the Aldh1a2 blocker WIN18446 reduced organoid formation and widespread inhibition of the Aldh1a family with 673A nearly eliminated organoid production (**Figure 3C**). Similar to control BRFL samples, D12 aggregates treated with AGN193109 produced OVs positive for the otic markers Pax2 and Sox2 (**Figure 3D-E, G-H**).

**Figure 3.**
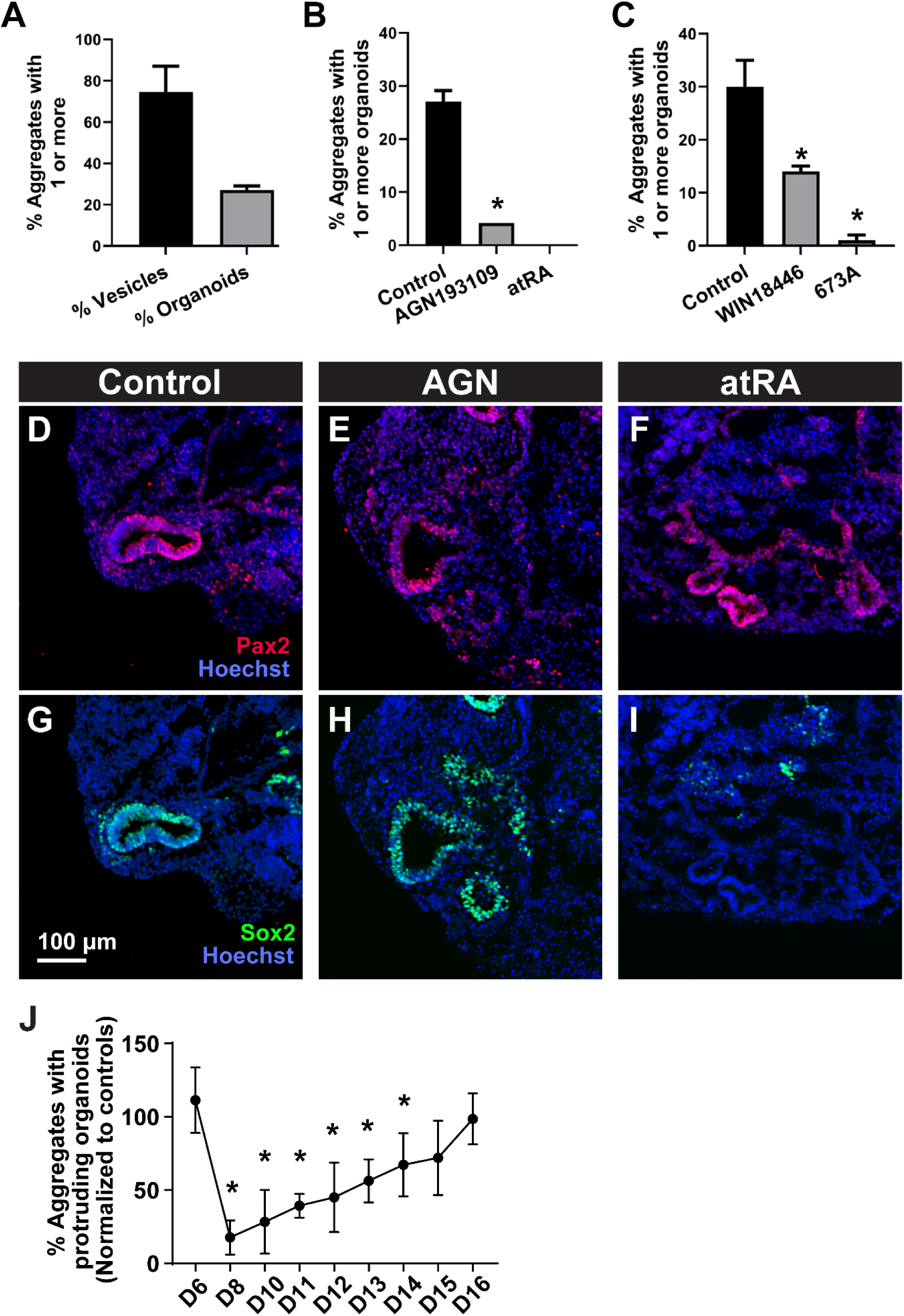
RA modulation of organoid efficiency. (A) The percentage of aggregates with 1 or more visible OVs at D12 or protruding organoids at D20 is shown for untreated control BRFL cultures. The efficiency of producing aggregates by D20 with protruding organoids is compared for (B) vehicle control cultures and those treated with the pan-RAR reverse agonist AGN193109 or 500 nM atRA and (C) vehicle control cultures and those treated with Aldh1a inhibitors WIN18446 or 673A. * *P*<0.05 in unpaired t-tests compared to control preparations. (D-I) Expression of OV markers Pax2 (red) and Sox2 (green), counterstained with Hoechst (blue) in cryosections from D12 aggregates, representative of over 20 aggregates per condition in 3 independent preparations. (J) A critical window of RA-sensitivity extends from D8 to D14, a time when otic placode and vesicle intermediates are being produced. The efficiency of producing aggregates by D20 with protruding organoids is shown for cultures treated with AGN193109 for 24 hours on the day indicated, D6-D16. Data were normalized to average control values. One-way ANOVA revealed a significant effect over time (*P*<0.0001) with * *P*<0.05 in post-hoc pairwise comparisons to control preparations. (A-C, J) mean ± one standard deviation with N = 3-5 independent samples per condition and 40-50 aggregates per sample. Scale: (D-I) 100 μm.

However, the addition of atRA appeared to eliminate Sox2 expression in the vesicles (**Figure 3F, I**), suggesting one possible mechanism for failed morphogenesis in this condition^40^. Additionally, in another similarity with *in vivo* otocyst development^15^, the influence of RA on organoid production in the BRFL cultures occurred in a critical developmental time window. We added AGN193109 to cultures for 24 hours, on D6 through D16 and compared the organoid production on D20 to vehicle controls (**Figure 3J**). Inhibition of RA activity on D6 had no impact on organoid-genesis. However, two days later at a time when the otic placodal domain begins to form, this manipulation led to an 80% reduction in cyst formation. Application of the inhibitor on subsequent days had steadily less impact until D15 when production resembled control levels.

### RA responsiveness in derived OVs is variable and asymmetric

Given the sensitivity of native and stem cell-derived OVs to RA manipulation, we hypothesized that variable RA activity may limit organoid efficiency. Mice with a RA reporter transgene have been used previously to reveal a transient wave of RA activity in mouse otic placode and OVs from E8.75 to E9.5^15^. These mutants carry random insertions of a transgene composed of three copies of the RA response element (RARE) from the RARβ receptor upstream of *lacZ*. To develop a RARE-lacZ mouse ESC line, we crossed Tg(RARE-Hspa1b/lacZ)12JRT mice males with wild-type C57BL/6J females. Hemizygous animals (RARE*^lacz/+^*) showed slightly elevated hearing thresholds compared to wild-type littermates, possibly due to the greater contribution of the CD1 background—which exhibits age-related hearing loss^41^— present in the RARE-lacZ mutants (**Figure 4A**). These mice showed normal cochlear morphology out to 4-months-of-age (**Figure 4B_1_**) and were able to maintain the ability to report on RA activity in early postnatal cochlea (P6) by X-gal staining (**Figure 4B_2_**). Over twenty E3.5 blastocysts from the F1 cross were examined for presence of the *lacZ* transgene, and line A6 was selected for analysis (**Figure 4C**). The RARE-lacZ ESCs showed a log-linear relationship between X-gal intensity and atRA dose, confirming their ability to sensitively report RA activity (**Figure 4D-E**). The RARE-lacZ ESCs were optimized for the organoid protocol, producing OVs, organoid cysts, and sensory hair cells (**Figure 4F-H**).

**Figure 4.**
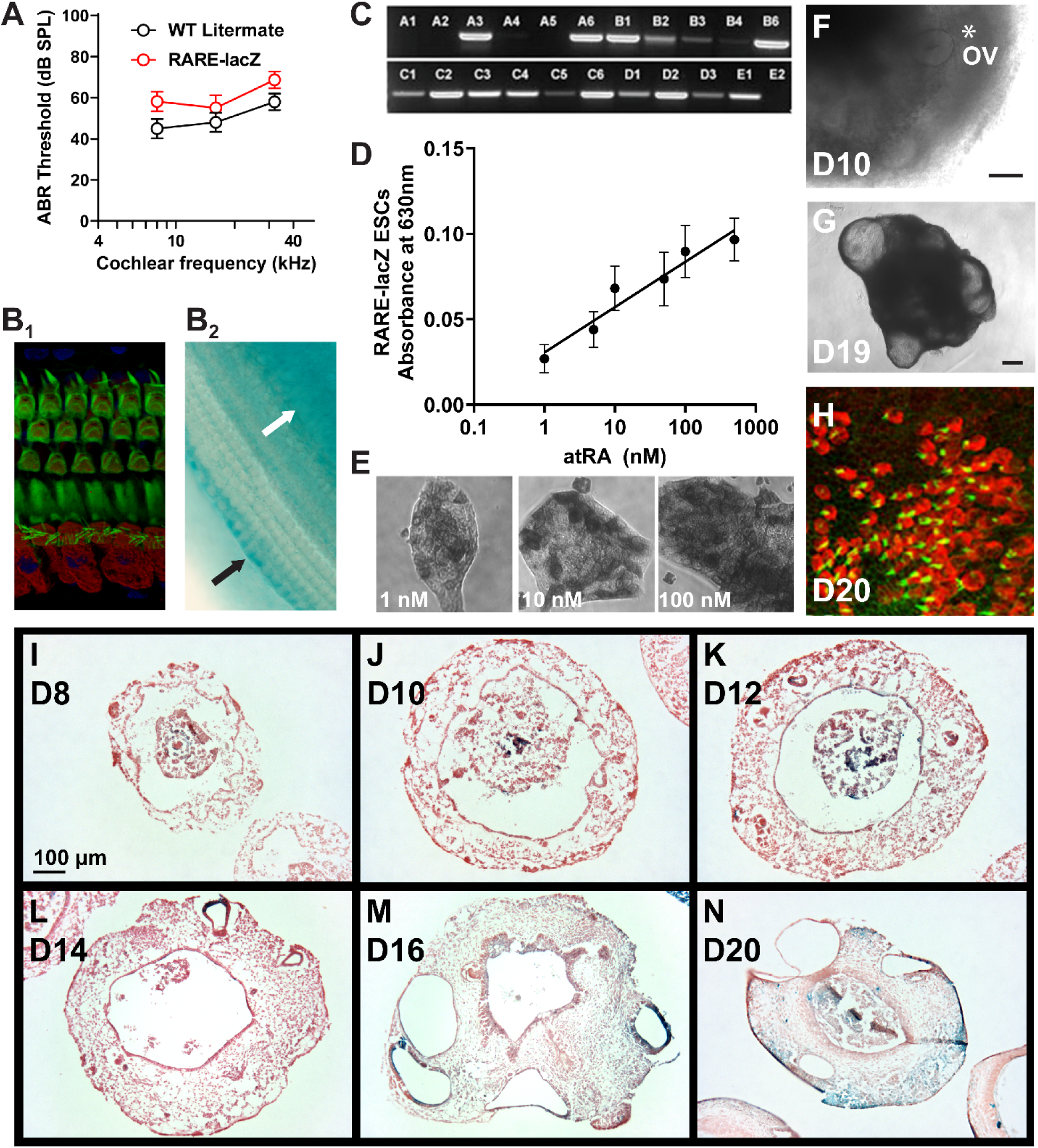
Development of RARE-lacZ reporter ESCs and inner ear organoids. (A) Auditory brainstem response thresholds for 4-week-old RARE-lacZ/+ mice and wild-type (WT) littermate controls (*P*<0.05, one-way ANOVA across genotype). Wholemounts of middle cochlear turns from RARE-lacZ/+ mice were stained by (B_1_) immunofluorescence with MyoVIIa (red) and phalloidin (green) or (B_2_) X-gal; arrows indicate positive Xgal stain in supporting cells bordering the OHCs (black arrow) and in the spiral limbus (white arrow). (C) Genotyping PCR for lacZ in 22 mouse ESC lines from RARE-lacZ/+ blastocysts. Clone A6 was chosen for follow-up experiments. (D) Absorbance from X-gal staining of line A6 exposed to a 24-hour atRA dose at various concentrations. Log-linear regression fit correlation coefficient (R^2^) was 0.94. (E) Increasing RA dose increased X-gal signal and the number of cells responding. The RARE-lacZ/+ ESCs were able to generate (F) OVs marked with asterisk and (G) organoids with (H) MyoVIIa-red and phalloidin-green labelled hair cells. (I-N) X-gal stain (blue) along with FastRed counterstain in representative thin sections are shown at indicated culture time points. Images are representative of 2 independent cultures and more than 10 independent aggregates per time point. Scales: (F-G, I-N) 100 μm

Whole aggregates from RARE-lacZ ESCs in the organoid protocol showed positive X-gal staining as early as culture D9, which became faint and more restricted to OVs at later time points (**see Figure S3A-C**). Localization of lacZ expression was more readily apparent in cryosections stained with X-gal, though there appeared to be some loss of signal compared to the whole aggregate preparations. At D8 and D10, RA activity was associated with the aggregate core, distinct from the surrounding placodal epithelium (**Figure 4I-J**). By D12 and following through D20, RA activity could be found within the OVs and organoids, but the labeling was highly variable between neighboring vesicles/organoids within the same aggregate and was non-uniform around their perimeters (**Figure 4K-N)**.

Since inhibition of RA signaling indicated a critical window of RA activity around the time of placode formation on D8, we tested whether sensitivity to excess RA was also maximal at this time point. The addition of 500 nM atRA on D8 increased lacZ expression intensely in whole aggregates at D9 (**see Figure S3A-C**), and this was largely due to intense staining of the otic-epibranchial placodal domain (**Figure 5A-B**). The impact of excess atRA was reduced when applied at later time points, consistent with a critical developmental window of RA sensitivity as the otic placode is formed (**see Figure S3A-C**).

**Figure 5.**
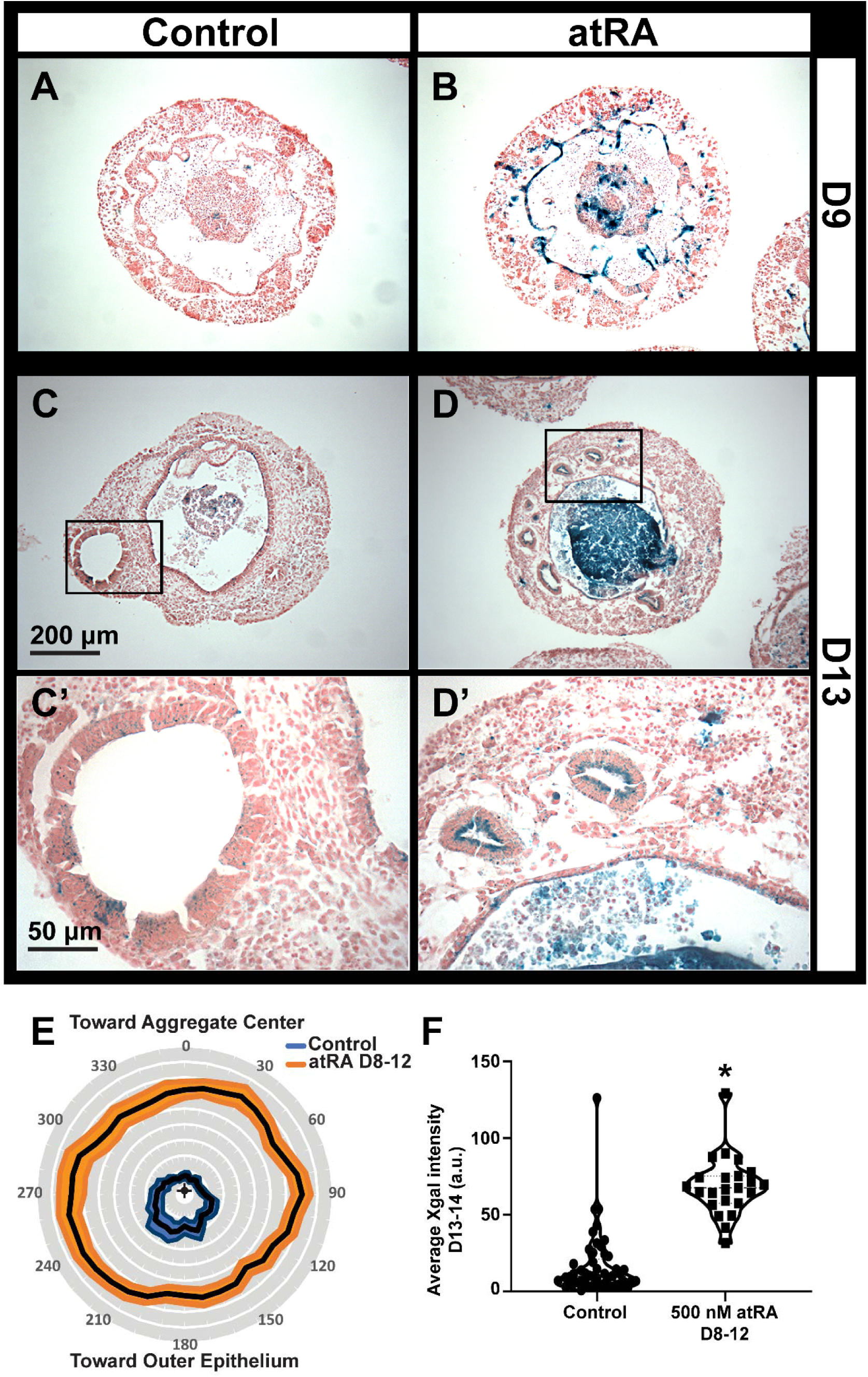
Asymmetry in RA responsiveness and impact of exogenous RA on RARE-lacZ reporter expression. Representative images from (A, B) D9 and (C, D) D13 preparations are shown for X-gal (blue) and FastRed stained sections from (A, C) control preparations and (B, D) those treated with 500 nM atRA from D8 to D12. Boxed insets in panels C and D are shown at higher magnification in C’ and D’, respectively. (E) Mean X-gal intensity for control (N=49) and atRA-treated vesicles (N=23) plotted on polar coordinates relative to the aggregate center. The mean is indicated by the center solid black line with one standard error of the mean shown in filled areas around the mean. One-way ANOVAs for each condition revealed a significant difference in intensity with respect to polar orientation for control samples (*P*<0.01) but not for those treated with RA (*P*>0.05). (F) The average X-gal intensity across each vesicle cross-section is shown by dot-plot with significance tested by unpaired t-test, \**P* < 0.0001. Scales: (A-D) 200 μm, (C’, D’) 50 μm

From these X-gal preparations, we also noted that the OV staining was non-uniform around the epithelium and assessed whether RA reactivity was polarized (i.e., asymmetric), potentially reflecting local signaling events. We stained BRFL cultures at D13 and D14—when X-gal intensity was greatest (**Figure 4L**)—to further examine the asymmetry and variability in RA response. Control cultures often exhibited asymmetric stains, with the more intense label occurring in regions away from the aggregate center, facing the outer edge of the aggregate (**Figure 5C,C’**). After treating with 500 nM atRA from D8-D12, the stain was more intense and more uniform around the cells lining the lumen of the OV (**Figure 5D,D’**). The perimeter of the OVs was also smaller in atRA-treated cultures than controls, 284.9±16.5 µm and 164.7±13.6 µm, respectively (mean ± one standard error of the mean; *P*<0.0001). We measured the X-gal intensity around the perimeter of the vesicles and transformed the data to polar coordinates (**see Figure S4A-B**). The average intensity for control vesicles was asymmetric, whereas atRA treatment caused more intense and uniform staining (**Figure 5E**). The overall X-gal intensity for each cross section was also compared. Control vesicles were highly variable, with some exhibiting intensity levels similar to the atRA-treated vesicles (**Figure 5F**), which corresponds to conditions that would arrest organoid-genesis (as shown previously in **Figure 3B**).

### Systematic variations in RA level regulates organoid efficiency and sensory-nonsensory fate

To gain control over RA activity, we sought to suppress endogenous RA synthesis and add atRA in various doses. Noticing some toxicity due to the commercial 673A pan-Aldh1a family inhibitor (e.g., smaller growth and loss of some epithelial integrity in the outer margins of the aggregates), we turned to custom Aldh1a inhibitors with greater specificity and potency (Compound 69 in Huddle et al., 2018 synthesized by the UM Vahlteich Medicinal Chemistry Core as Compound CCG-263646, or 646 for short^42^). Inhibitor 646 was applied during placode and OV formation from D7 to D12 while atRA was added from D8 to D12 at doses from 0 to 500 nM. Exemplary images of D20 aggregates show a non-monotonic dose-dependency on production of protruding cystic organoids (**Figure 6A**). Cryosection staining of these aggregates for hair cell markers revealed internal hair cell-containing organoids in the absence of RA though we never saw protruding cysts in these preparations, low levels produced large numbers of hair cells, and high levels of atRA produced limited or no sensory cells (**Figure 6B**). The efficiency of organoid production was quantified over several independent preparations and intermediate levels of atRA could produce up to 2.5-fold more aggregates with one or more protruding cysts (**Figure 6C**). Hair cell density was dose dependent with the highest density at low atRA levels, though cyst area much like organoid efficiency was non-monotonic and peaked at moderate atRA levels (**Figure 6D-E**).

**Figure 6.**
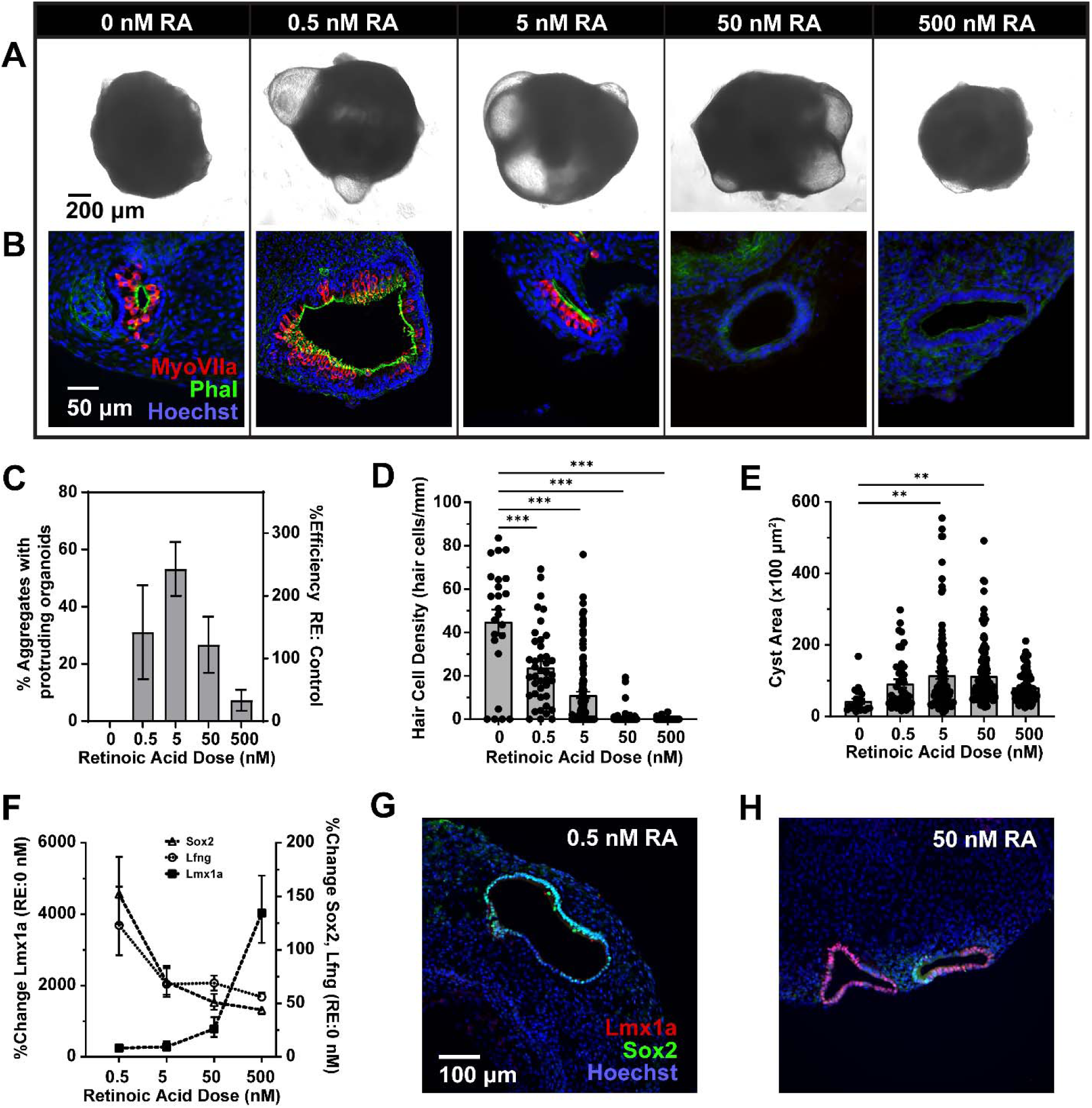
Effects of controlled exposure to exogenous atRA on sensory-nonsensory cell fate in inner ear organoids. Endogenous RA production was suppressed with the pan-Aldh1a inhibitor 646 applied from D7-D12 while cultures were treated with various concentrations of exogenous atRA on D8-D12. (A) Example aggregates are shown for each RA condition alongside (B) representative immunofluorescence images of cryosections stained for MyoVIIa (red), phalloidin (green), and Hoechst (blue). (C) Efficiency of the production of protruding cyst-like organoids, shown as both the % aggregates containing one or more organoids and the average percentage relative to control (untreated) cultures. (D) Hair cell density and (E) organoid cyst area is shown for thin sections from D20 aggregates under the various RA conditions. Hair cell number was quantified as density, normalized by the perimeter of the cyst within the section. Dots indicate individual data points on top of bar graphs of mean ± one standard error of the mean. (F) Quantitative PCR data is shown for the % change in prosensory markers *Lfng* and *Sox2* and the nonsensory marker *Lmx1a* relative to the 0 RA condition (646 in the absence of atRA). Thin sections from D20 organoids are shown stained for the sensory domain marker Sox2 (green) and nonsensory marker Lmx1a (red), counterstained with Hoechst under low-dose RA (G, 646+0.5 nM atRA) and high-dose RA (H, 646+50 nM atRA). (C-F) mean ± standard deviation with N = 3-6 independent samples per condition. * *P*<0.05, \*\**P*<0.01, \*\*\**P*<0.001 in post-hoc pairwise comparisons compared to control in (C) and 0 RA condition in (D-E). Scales: (A) 200 μm, (B) 50 μm, (G-H) 100 μm.

Increasing atRA dose resulted in decreasing sensory (i.e., *Sox2* and *Lfng*) and increasing nonsensory (i.e., *Lmx1a*) gene expression in whole aggregates from D12 cultures (**Figure 6F**). These relationships were statistically reliable using one-way ANOVA on the ΔCt values for each gene (*Sox2 P*<0.01, *Lfng P*<0.05, *Lmx1a P*<0.0001). These effects were validated at the protein level with immunofluorescence on D12 cryosections using antibodies to Lmx1a and Sox2 (**Figure 6G-H**). At low levels of RA, only Sox2-positive prosensory domains were found in D12 OVs, while at higher doses the OVs showed increasing amounts of Lmx1a nonsensory domains. By controlling RA exposure, organoid fate could be systematically shifted from sensory to non-sensory, but the size and efficiency in producing large organoid cysts appeared to occur at intermediate RA levels.

## DISCUSSION

In this study, parallels between early stages of IEO formation and mouse otic development were explored with an eye toward identifying cues that could further tailor cell type specification and modulate culture efficiency. The efficiency of each step in differentiation depends upon the efficiency of each preceding step. We modulated two factors that bookend the cue-driven phase of organoid culture paradigms: the first (TGFβ) modulating germ layer identities and the second (RA) modulating axial patterning of otic intermediates. In the first case, the suppression of mesendoderm by TGFβ inhibition at initial stages is pivotal in the production of surface ectoderm from which placodal domains, OVs, and IEOs will emerge over the course of the next several weeks *in vitro*^8^. Given the fundamental role TGFβ signaling plays in embryos and in early differentiation of stem cells, it was a logical target to begin optimizing the IEO method to reach its full potential.

In our cultures, TGFβ inhibition was necessary for vesicles to mature into hair cell-containing organoids. However, Sox2-positive OVs were observed in the maturation phase with or without of TGFβ inhibition in the earlier ectodermal differentiation phase. Exclusion of SB431542 had no discernible impact on vesicle formation or character. This was surprising given that attenuation of TGFβ signaling by endogenous antagonists is thought to be necessary for formation of anterior ectoderm in developing mouse embryos^43–46^. The TGFβ pathway’s role in body axis patterning and germ layer specification is conserved across vertebrate and invertebrate species^47^. Likewise, its inhibition promotes an ectodermal lineage for mouse ESC aggregates on the path toward otic fate via the IEO protocol. One explanation is that our panel of OV markers may be too narrow or the immunostaining approach not sensitive enough to detect subtle changes in functionally significant processes for further otic differentiation. Regardless, the improved efficiency of vesicle formation per aggregate with RepSox and SIS3 supports these inhibitors as useful alternatives to SB431542 with RepSox becoming a key part of our organoid protocol in several other reports^3,48^.

Since aggregates without TGFβ inhibition ultimately failed to produce organoids, the impact of this inhibition was ostensibly delayed until after the vesicle stage. This delay suggests an epigenetic mechanism, which is a primary means of TGFβ influence over development, immunity, and regeneration in many cell contexts^49^. In development, epigenetic changes have been suggested to explain the delay between onset of competence factors and preplacodal markers, with most of gastrulation occurring in the interim^26^. Embryonic stem cells show global DNA demethylation so that progressive epigenetic changes may underlie commitment to lineage paths; in addition, epigenetic mechanisms may play an active role in fate decisions^50,51^. Investigating epigenetic changes in the derived vesicles not only could inform optimization of the IEO protocol by revealing targets for demethylation to re-open blocked lineage paths, for instance, but could also provide new insights into mechanisms involved in early embryonic development.

Principal component analysis and hierarchical clustering of the RNASeq data revealed that, relative to untreated vesicles, SB431542-treated OVs represented a shift towards the native E10.5 transcriptome. In the expression levels of some genes, however, SB431542-treated vesicles were still more “derived” than “native-like,” as evidenced by the proximity of BSFL and BFL clusters compared to E10.5. Achieving a more native-like vesicle through additional optimization of the IEO protocol is a goal of ongoing studies. Focusing on the 4,543 genes differentially expressed between native and derived vesicles and unaffected by TGFβ inhibition may reveal new targets for additional exogenous cues. Our analysis of the DEGs at this intersection suggested at least two immediate options to pursue. One is to further optimize the type, concentration, and duration of TGFβ inhibition, since even BSFL OVs show a persistent upregulation of genes in non-ectodermal lineages compared with native tissue. The other factor highlighted by our study was associated with RA signaling.

Retinoic acid is implicated in patterning the anterior-posterior axis of the OV^12^. The otic placode is exposed to opposing gradients of RA synthesizing and degrading enzymes as the placode invaginates to form the otic cup ^15^. By the time the OV has fully formed, the epithelium is no longer responsive to RA until later stages of organ maturation^52,53^, yet RA is crucial to establish anterior (e.g., Lfng and NeuroD) and posterior (e.g., Tbx1) marker domains within the OV^15^. During a critical developmental window, perturbations in RA—creating excess or deficiency—results in OV dysmorphogenesis^17^, which our data replicate in organoid cultures. Investigations into the mechanisms of RA control over OV morphogenesis has led investigators to suggest that deficits due to excess or deficient RA are largely due to changes in Fgf3/10 in periotic mesenchyme and ultimately in the downstream targets Dlx5/6 expressed within the otocyst^17^. Our data adds differential effects on Sox2 as another key player in teratogenicity from excess RA. The impact of RA signaling on a cell is context dependent; that is, RA has been shown to increase Sox2 and other factors to drive cell proliferation or decrease Sox2 and other factors to guide differentiation^54^. Sox2 is essential for normal inner ear development; early deletion in the otocyst results in the loss of sensory domain formation and severe dysmorphogenesis of the inner ear^40^. In the organoid cultures, excessive atRA eliminated Sox2 in the derived OV and expanded Lmx1a-positive nonsensory domains, leading to the loss of large protruding cysts and differentiation of sensory hair cells.

Context dependent roles for RA have been reported in a variety of tissues and for many target genes; RA influence over the downstream effector Tbx1 is one example. In the normal mouse posterior OV, RA appears to promote the expression of Tbx1^15^, and this relationship may be generalizable to other placodes^55^. In contrast, RA and Tbx1 are mutually repressive in non-placodal tissues^56,57^. Our data, surprisingly, fit with this non-placodal model, where gene expression data and X-gal staining results suggested persistent RA activity in the organoid cultures though *Tbx1* was downregulated compared to E10.5 tissues. As further evidence, *Otx1* expression, which is downstream of *Tbx1*^58^, was also higher in native than derived vesicles. It is also possible that our isolation of OVs from the larger, cellularly complex aggregate may have resulted in unintentional contributions from mesenchyme. For example, Hox genes, which are primarily expressed in paraxial mesoderm^37^, were expressed higher in derived than native samples. A single-cell atlas may be required to further tease apart the relationships between RA and downstream targets in different otic tissues and at different developmental stages.

Our data indicated that RA sensitivity in mouse IEO cultures, when treated with exogenous atRA, was greatest around the time of placode formation and evagination of the otic cup. This timing is similar to the RA responsivity in normal inner ear development from E8.75 to E9.5^15^. However, in control cultures, RA responsivity was greatest during OV formation and persisted in many OVs throughout organoid-genesis. This high, persistent RA likely contributes to differences in the efficiency, morphology, and fate of the organoids. Physiological evidence of aberrant endogenous RA signaling in the culture paradigms included (1) reductions in organoid efficiency upon blockade of RA receptors, (2) similar reductions in efficiency upon blockade of endogenous RA synthesis, (3) a dose-dependent effect on efficiency when blocking select synthesis enzymes compared to all Aldh1a family members, (4) RARE-lacZ responsivity localized to otic intermediates, and (5) variable response in RARE-lacZ intensity with the highest responsivity associated with conditions that fail to produce organoids. The source of the endogenous retinoids is unclear, but there remain some components of the culture paradigm that are ill-defined, such as the animal-derived artificial extracellular matrix Matrigel. Most importantly, by suppressing endogenous RA synthesis and applying exogenous levels of atRA, we were able to increase organoid efficiency and systematically vary fate decisions between sensory and nonsensory domains. The direct targets of RA in the ear remain unknown, but this culture system could open new avenues for exploring dose-dependent transcriptional regulation by RA at various stages of inner ear development. In addition, several investigators have suggested that RA can induce re-entry of mature inner ear supporting cells into the cell cycle as a means for hair cell regeneration^52,59^. Our culture system could be used to explore this by systematically varying RA concentrations at later stages of organoid development, specifically after D16 beyond the critical window of RA impact on organoid production.

In addition to changes in efficiency and cell fate, RA impacted organoid morphology. There are at least two major morphological types of mouse IEOs: those that are fully embedded in the aggregate and large protruding cysts^7^. Embedded organoids are smaller and tend to have a thicker epithelium, while protruding organoids are large with substantial thinning of the epithelium. To simplify determinations of organoid production efficiency, we tend to quantify the number of aggregates with one or more protruding organoids, as these are readily identifiable under a cell culture microscope. Thus, observations of dose-dependent effects on organoid efficiency largely reflected changes in the generation of protruding cysts. However, in thin sections, we found embedded, internal organoids at all RA doses. Nothing is currently known about the composition and homeostasis of the luminal fluid of embedded or protruding organoids. However, it seems inescapable that the composition of ion channels and transporters are unique between these two types of cysts. Changes in ion flux across the epithelia would be accompanied by changes in water movement, hydrostatic pressure, and bulk volume. The resulting changes in the mechanobiology of the surrounding epithelium would then be expected to feedback onto cell fate decisions, as appears to be the case in other organoid systems^60^.

In summary, we have demonstrated that RA and TGFβ—two morphogens that show considerable crosstalk in development^61^ and play major roles in establishing the anterior-posterior body axis early in embryogenesis^43,62^—have a profound impact on the production of IEOs. Future investigations should attempt to tease apart the intersections of these pathways in organoid-genesis.

Understanding how these, and other, signaling events influence efficiency and cell fate at each stage will be especially beneficial to inner ear research as it may lead to generating large quantities of hair cells for *in vitro* studies and for developing *in vivo* hair cell replacement therapy.

## Acknowledgements

This work was supported by grants from the Veterans Affairs (I01 RX003400 to R.K.D.) and NIH NIDCD (T32 DC000011 for S.A.S). The authors are thankful for the assistance of Drs. Stacy Schaefer and Joerg Waldhaus for some specimen and comments on early drafts of the manuscript. The authors also acknowledge Matthew Adams for initial assistance with evaluating the RARE-lacZ ESCs.

## Author Contributions

R.K.D. conceived and performed experiments, wrote the manuscript, and secured funding; L.L., M.M., A.W, R.D., and L.C. conducted the experiments.

## Lead Contact

Requests for resources and additional information should be directed to the lead contact, R. Keith Duncan.

## Resource Availability

Mouse cell lines generated in this study are available from the lead contact with a completed materials transfer agreement. Raw data from the RNASeq analysis were deposited at GEO and are publicly available with accession number GSE285576.

## Supplemental Information

Document S1. Figures S1-S4 and Tables S1-S2

Table S3. Excel file containing additional data of DEGs in pairwise comparisons too large to fit in a PDF

## STAR Methods

### Key resources table

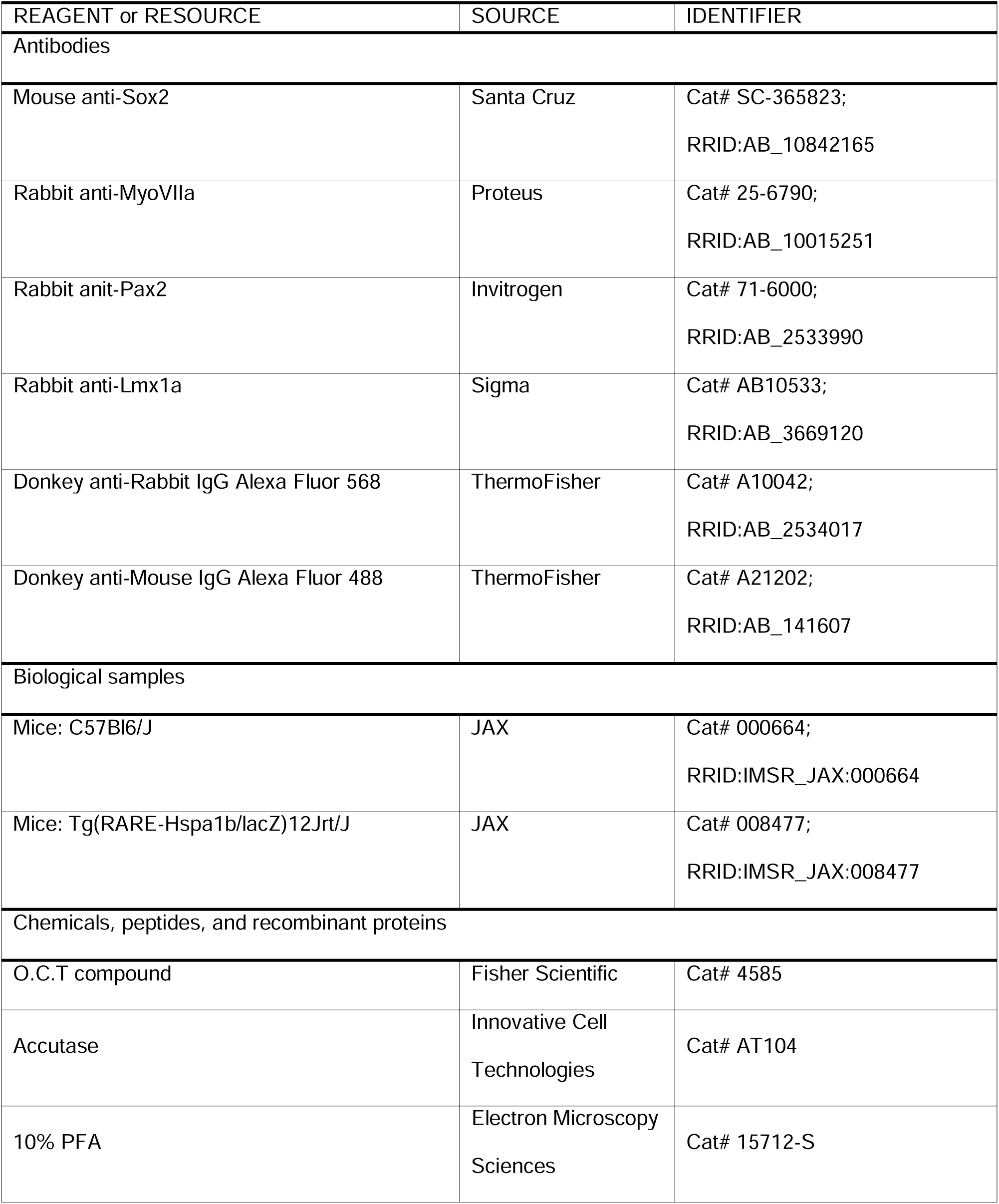
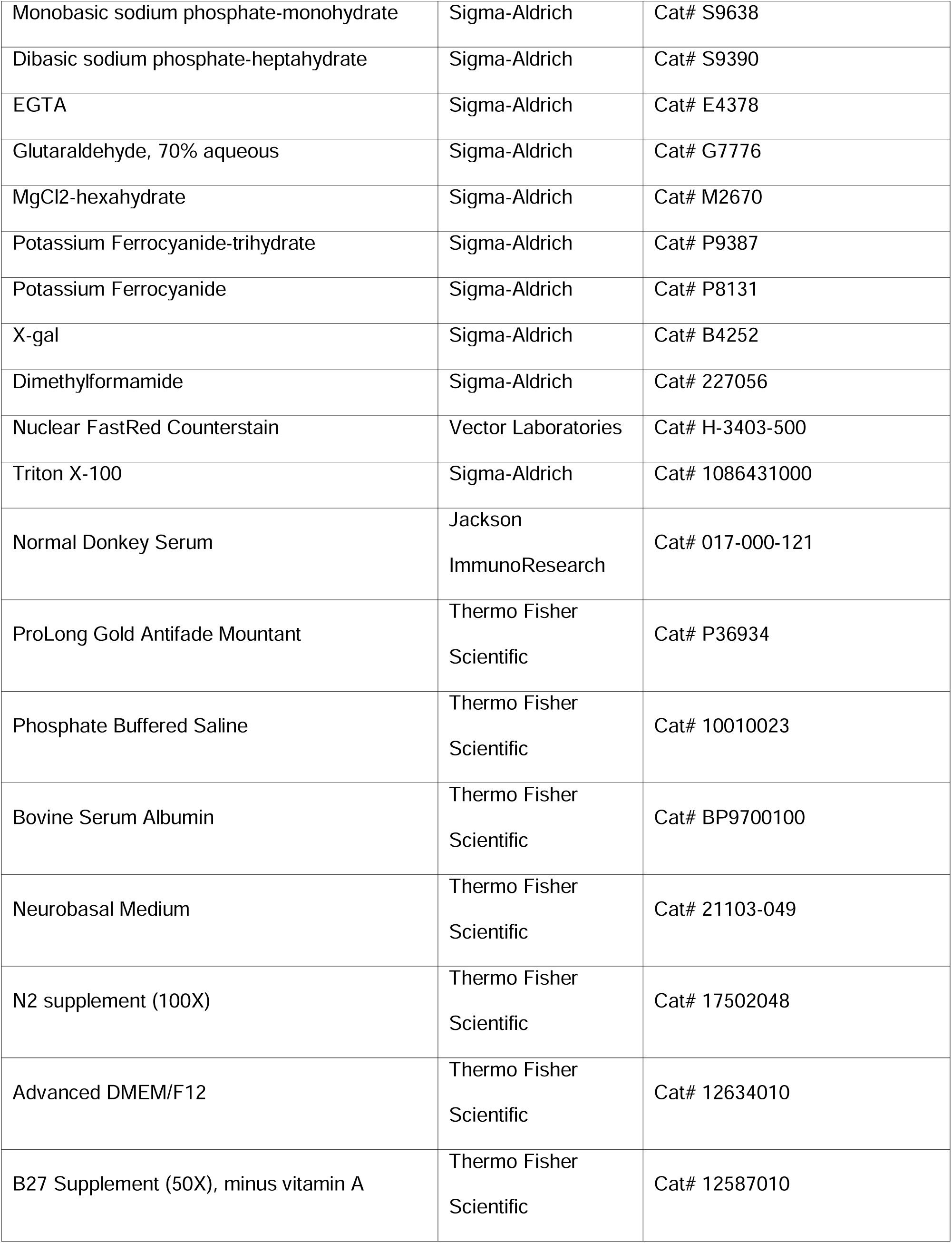
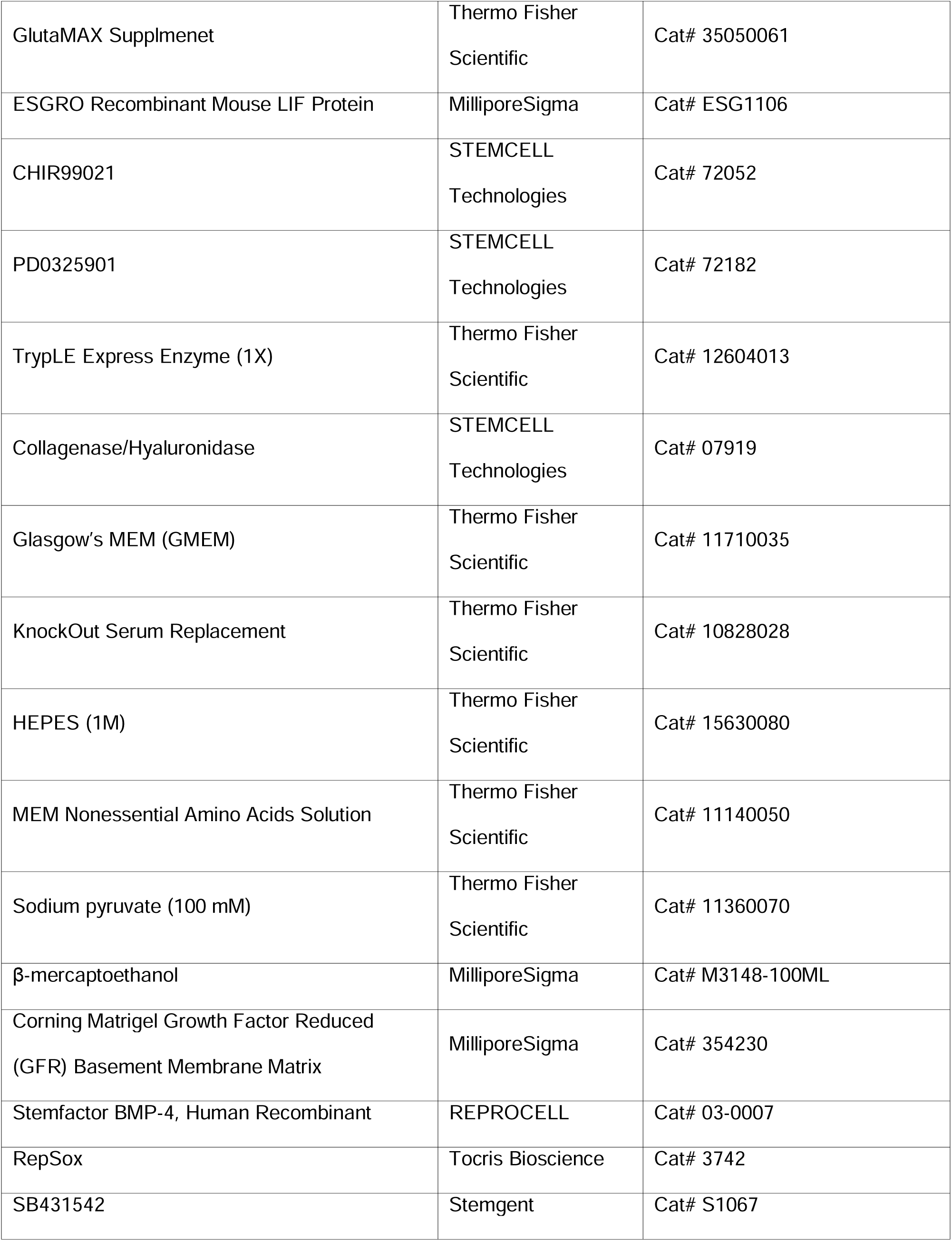
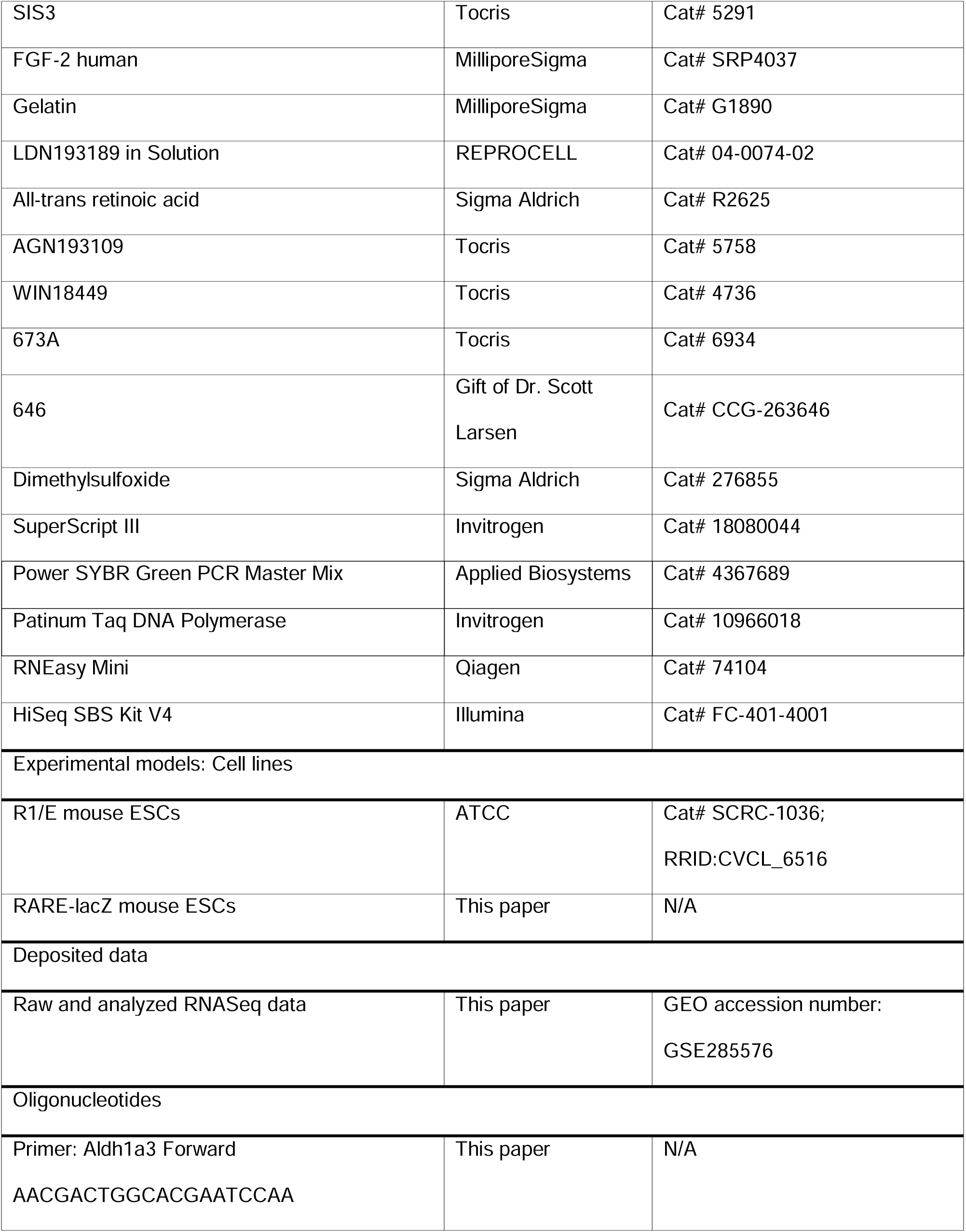
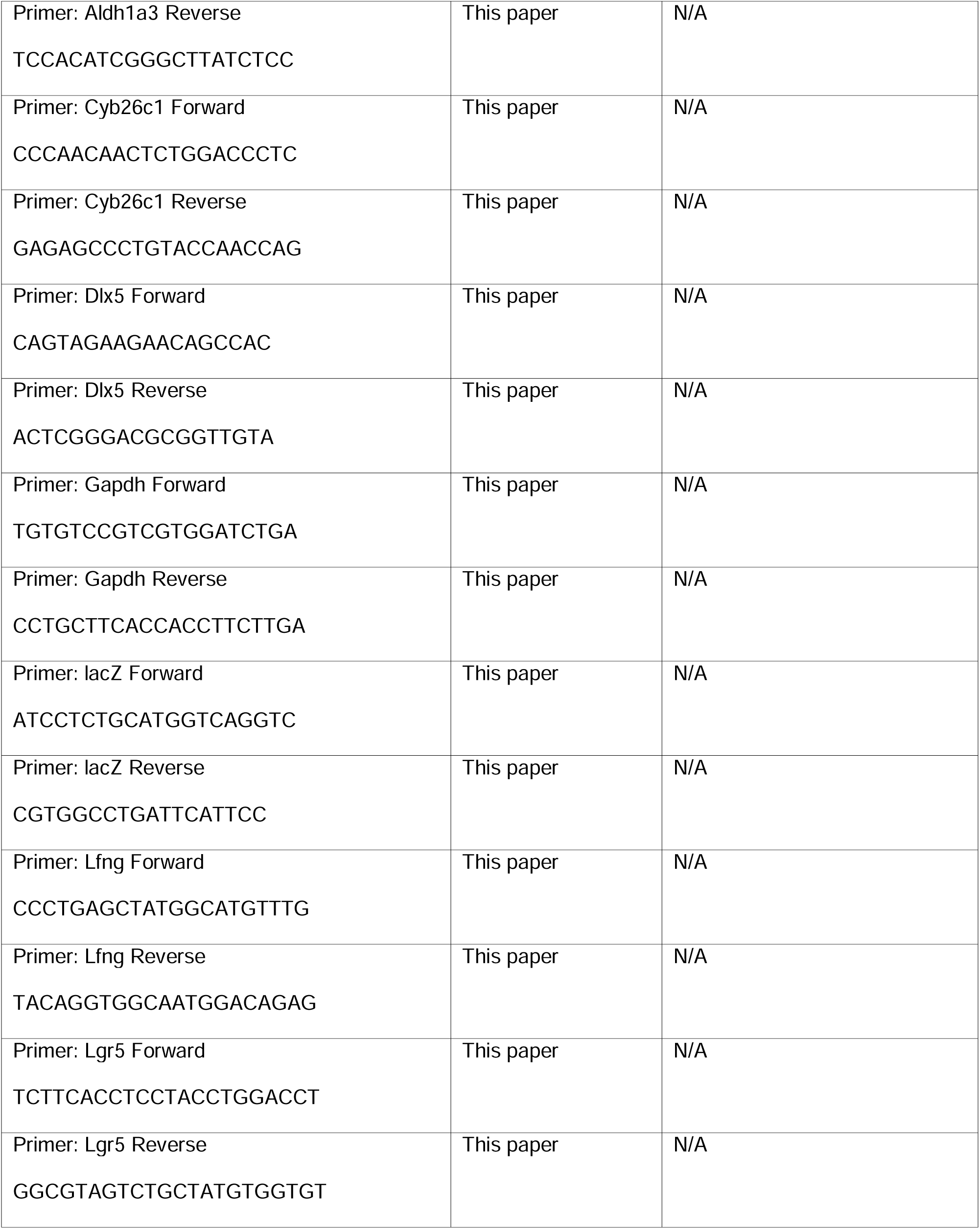
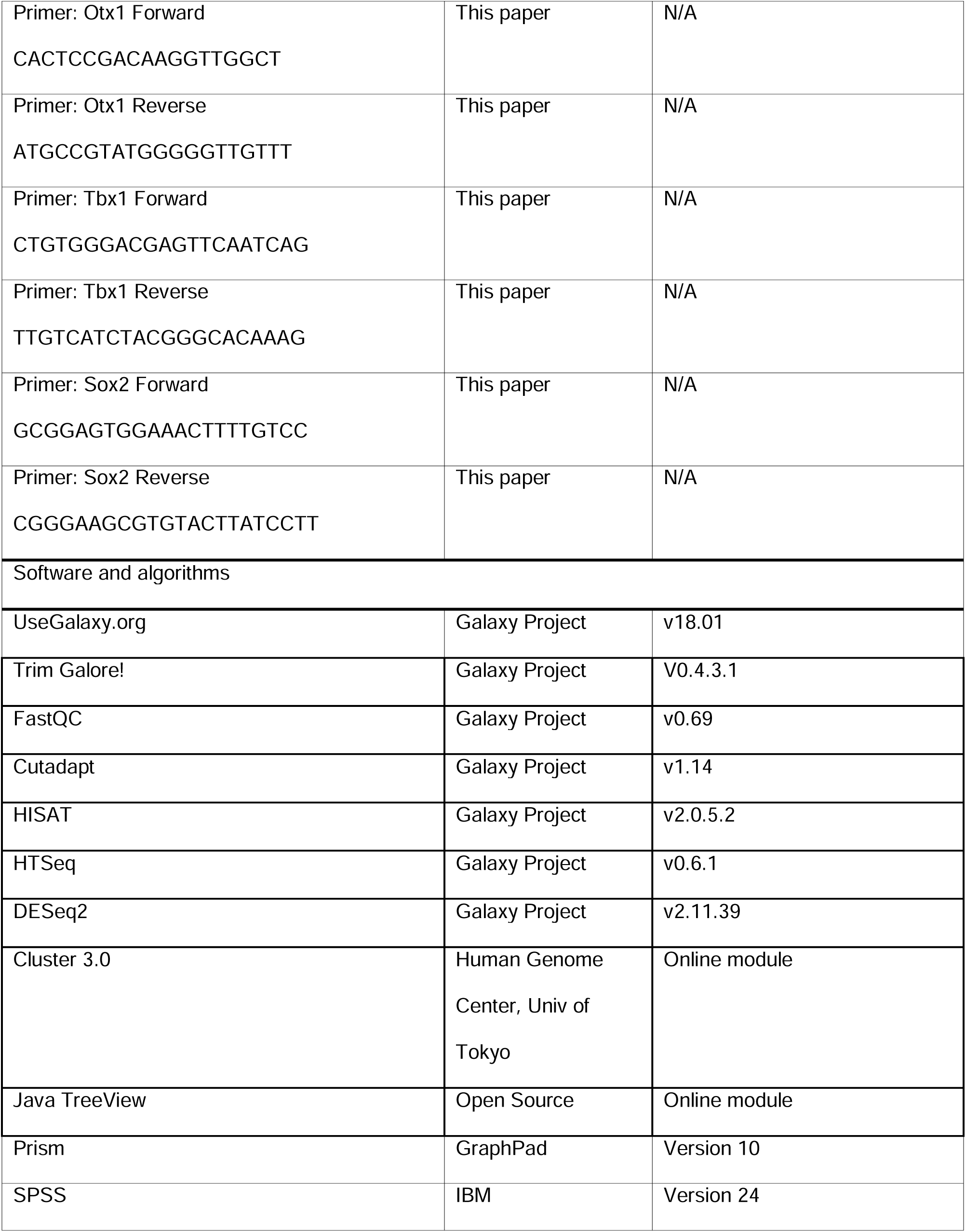
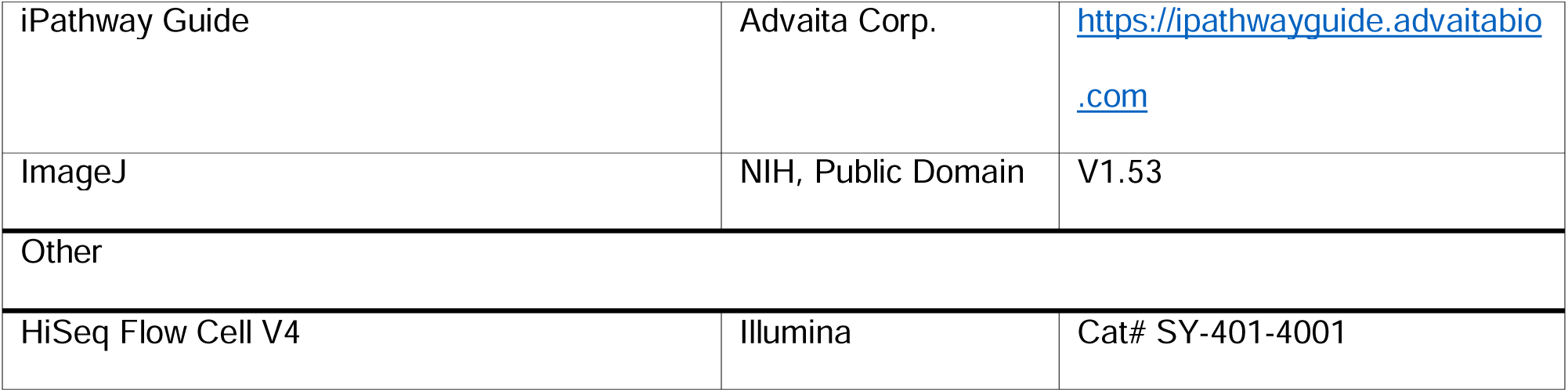

### Experimental models

The RARE-lacZ ESC line was established in collaboration with the University of Michigan Transgenic Animal Core, crossing RARE-lacZ (Tg(RARE-Hspa1b/lacZ)12Jrt) males with wild-type C57BL/6J females, purchased from Jackson Laboratory. All animal maintenance and experimental procedures were performed in accordance with NIH guidelines and were approved by the Institute Animal Care and Use Committee at the University of Michigan (Protocols PRO00008446 to R.K.D. and PRO00007879 to Dr. Thom Saunders in the UM Transgenic Animal Core Facility). The mutant RARE-lacZ mice carry a transgene with three copies of RA response elements (RARE) upstream of lacZ^63^. Embryos for establishing ESC lines were harvested from timed pregnant dams with the pug date referencing E0.5. Blastocysts from E3.5 embryos were expanded on feeder layers of mitotically inactivated mouse embryonic fibroblasts before adapting to feeder-free culture conditions for maintaining the ESCs. In addition, the study utilized R1/E mouse ESCs originally purchased from ATCC (SCRC-1036). Frozen stocks of mouse ESCs were maintained in vapor phase of liquid nitrogen storage.

### Mouse ESC cultures

For stem cell maintenance, colonies were cultured in feeder-free conditions on 0.1% gelatin in 2i culture media consisting of a 1:1 mixture of Advanced DMEM/F-12 and Neurobasal, 0.5X B-27 (without vitamin A), 1X N-2 supplement, and 1X GlutaMAX supplemented with 1000 U/mL LIF, PD0325901 (1 μM), and CHIR99021 (3 μM). Colonies were dissociated to single cells with TrypLE Express (Gibco) for maintenance and for producing aggregates.

### Differentiation protocol

Mouse ESC aggregates were cultured according to the previously described inner ear organoid protocol [1,2,15]. On D0, following dissociation of colonies, cells were reaggregated in round-bottom 96-well Nunclon Sphera Microplates (Thermo Scientific). Cells were seeded at a density of 3,000 cells in 100 μL ectodermal differentiation medium per well. The medium was composed of GMEM, 1.5% KnockOut serum replacement (KSR), 15 mM HEPES, 1X non-essential amino acids, 1 mM sodium pyruvate, and 0.1 mM β-mercaptoethanol. On D1, growth factor reduced (GFR) Matrigel was applied at a final concentration of 2% in ectodermal differentiation medium by replacing half the volume in each well. On D3, 10 ng/mL BMP4 was added in each well, with or without TGFβ inhibition. Inhibitors tested included SB431542 (1 μM), SIS3 (3 μM), and RepSox (1 μM). On D4.5, 1 μM LDN193189 and 100 ng/mL FGF2 were added. To begin the maturation phase on D8, aggregates were transferred into maturation medium consisting of Advanced DMEM/F-12, 1X N-2 supplement, 15 mM HEPES, and 1X GlutaMAX, with 1% GFR Matrigel and 3 μM CHIR99021. Half the volume of media was exchanged daily beginning on D10. Aggregates were monitored for vesicle formation by D12 and for organoid formation by D20-22.

### Isolation of otic vesicles

For isolating E10.5 otic vesicles from mice, pregnant C57BL/6 dams were euthanized, and uterine horns were removed and placed into PBS on ice. Otic vesicles were harvested from embryos by creating a window in the epithelium adjacent to the second branchial arch using a scalpel blade. Vesicles were then teased away from periotic mesenchyme using fine forceps. Isolated vesicles were transferred to Buffer RLT in RNase-free tubes on ice and then frozen quickly on dry ice before storage at -80°C. Separate pregnant dams were used for each of the 4 biological repeats; each repeat comprised 2-6 OVs from 1-3 embryos.

For isolating D12 otic vesicles from organoid cultures, R1/E aggregates were incubated in 1X collagenase/hyaluronidase in 35-mm Nunclon Sphera dishes placed at 37°C in a humidified 5% CO_2_ culture incubator for 45 minutes. At 15-minute intervals, aggregates were triturated gently with cut 1-mL pipette tips to encourage gradual disruption and aid diffusion of the enzymes. After 45 minutes, an uncut tip was used to fully dissociate aggregates to a mixture of single cells, residual clumps, and intact vesicles. The mixture was filtered through a 40-μm cell strainer, which was then inverted and washed with DMEM/F-12 to retrieve vesicles into a fresh 35-mm Nunclon Sphera dish. In some cases, vesicle number was quantified by counting isolates and normalizing to the number of aggregates in the sample. For other downstream applications, vesicles were collected in Buffer RLT (Qiagen) in RNase-free tubes on ice and then frozen quickly on dry ice before being transferred to -80°C until RNA extraction. Selection was limited to 20 minutes to avoid RNA degradation. At minimum, 30 vesicles were collected from each biological repeat, and 3-4 biological repeats were performed per condition.

### Quantification of vesicles and organoids efficiency

The rate of vesicle and organoid production was estimated under light microscopy. Vesicles were identified in the translucent margins of the aggregate on D10 to D12. Inclusion criterion included observation of fully intact spherical vesicles with a thickened epithelium and fluid-filled lumen. Organoids were identified as protruding of translucent cysts between D20 and D22, having a well-defined epithelial border. The rate of vesicle or organoid production was expressed as the percentage of aggregates with at least one object of interest meeting inclusion criteria. At least 32 aggregates were screened per condition per trial.

### RNA sequencing and analysis

Total RNA was extracted using RNeasy Mini Kits (Qiagen) and transferred to the UM Advanced Genomics Core for library preparation and sequencing. RNA input was assessed for quality and quantity using an Agilent Bioanalyzer 2100, Agilent 2200 TapeStation, and NanoDrop ND-1000 spectrophotometer (Thermo Scientific). The RIN scores ranged from 8-10. cDNA libraries were prepared from 100 ng total RNA per sample using a TruSeq RNA Sample Prep Kit v2 (Illumina). Library quality was confirmed by TapeStation and qPCR before sequencing. The Illumina HiSeq-2500 platform was used to perform V4 single end, 50 bp sequencing of libraries. Samples were sequenced in duplicate, with each sample loaded in two separate lanes. Fastq output files generated by bcl2fastq software v2.17 (Illumina) were uploaded to the Galaxy web platform (http://usegalaxy.org/).

Data analysis was performed using Galaxy v18.01^64^. Within Galaxy, the wrapper Trim Galore! v0.4.3.1 was used to assess the quality of base calls via FastQC v0.69^65^ and to trim and filter reads via Cutadapt v1.14. Trimming removed low-quality base calls (Phred < 20) before adapters, and then filtering removed short reads (<20 bp). Reads were aligned to the mm10 genome assembly with HISAT2 v2.0.5.2^66^. The resulting BAM files were merged to combine data from duplicate lanes. Data were converted to raw counts (reads per transcript) with the HTSeq v0.6.1 script htseq-count using Ensembl annotations^67,68^.

Raw counts were then normalized for differential expression analysis using DESeq2 v2.11.39^69^. Normalized counts from DESeq2 were processed using Cluster 3.0^70^ for preparation of heatmaps using Java TreeView^71^. Functional analysis based on Gene Ontology enrichment was performed on HTSeq counts within iPathwayGuide^72,73^, setting threshold P-values at < 0.05 adjusted for false discovery rate and Log_2_FC>0.6.

### Auditory brainstem response

Auditory brainstem responses (ABRs, the summed activity of auditory afferent pathways to short tone bursts) were performed on 1-month-old mice anesthetized with a mixture of ketamine (100 mg/kg, i.p.) and xylazine (20 mg/kg, i.p.). ABR acoustic stimuli were delivered through a closed acoustic system, consisting of two sound sources (CDMG15008-03A, CUI) and an electret condenser microphone (FG-23329-PO7, Knowles) as an in-dwelling probe microphone. Three needle electrodes were placed into the skin at the dorsal midline: one close to the neural crest, one behind the left pinna, and one at the base of the tail (ground). ABR potentials were evoked with 5 ms tone pips (0.5 ms rise-fall, with a cos^2^ envelope, at 40/s) delivered to the eardrum at 8, 16 and 32 kHz. The response was amplified (10,000X) and filtered (0.3–3 kHz) with an analog-to-digital board in a PC-based data-acquisition system. Sound pressure level was raised in 5 dB steps from 20 to 80 dB SPL. At each level, 1024 responses were averaged (with stimulus polarity alternated) after “artifact rejection” above 15 μV. ABR thresholds were determined as the lowest SPL were the first peak of the ABR waveform was still visible using ABR Analysis Software (Mass Eye and Ear, Boston, MA).

### Whole mount preparations, cryosections, and immunostaining

Wholemounts of cochlea from 6-day-old neonatal RARE-lacZ mice and organoid cysts were prepared for immunostaining to identify sensory hair cells. Animals were euthanized under anesthesia and the bony cochlear duct isolated into fixative (4% paraformaldehyde) for several hours at room temperature. The organ of Corti was microdissected from the cochlear duct and maintained in PBS. Whole organoid aggregates were fixed similarly, and large organoid cysts microdissected away from the larger spheroid. In some cases, aggregates were cryosectioned prior to immunostaining. After fixation, these specimens were preserved in 30% sucrose, embedded in OCT, and frozen on dry ice. OCT blocks were sections using a Leica 3050S cryostat at 10-12 µm thickness. Sections were dried overnight at room temperature prior to rehydration and downstream examining by immunofluorescence.

For immunostaining, cochlear wholemounts, dissected organoids, or cryosections were blocked in 10% normal donkey serum and permeabilized in 0.1% Triton X-100 before incubating overnight with primary antibodies in a 1:1 solution of PBS and blocking/permeabilization solution. Specimen were then incubated with Alexa Fluor secondary antibodies in PBS at room temperature for 1-2 hours. Hoechst 33242 was used for nuclear counterstaining. Alexa Fluor-conjugated phalloidin was paired with secondary antibodies in some cases, for the purpose of labeling actin-rich hair bundles. Each preparation was mounted with ProLong Gold Antifade Mountant prior to imaging.

### Preparation and imaging of X-gal-stained specimen

Cochlear wholemounts from RARE-lacZ hemizygous mice and whole RARE-lacZ aggregates from organoid cultures were fixed with 4% paraformaldehyde and 0.5% glutaraldehyde in 0.1M phosphate buffer for 1 hour at room temperature. In some cases, aggregates were cryopreserved as above and sectioned prior to X-gal staining. Intact specimens were permeabilized in wash buffer composed of 0.02% NP-40 and 0.01% sodium-deoxycholate in phosphate buffer with 2 mM MgCl_2_ to aid in penetration of the staining agents. Pre-warmed X-gal stain was prepared as 1 mg/ml X-gal in dimethylformamide in a buffer solution of 5 mM potassium ferrocyanide and 5 mM potassium ferricyanide. Preparations were incubated in this staining solution overnight in a humidified chamber at 37°C. After washing extensively, samples were counterstained in Nuclear Fast Red for 5 minutes at room temperature followed by PBS wash and imaging under phase contrast.

To quantify X-gal staining on cyrosectioned aggregates from D8 to D20, images were obtained from staggered sections approximately 200 µm apart to avoid analyzing the same object on different sections. Images were imported into ImageJ for analysis, including deconvolution to isolate the blue X-gal stain from Fast Red, followed by tracing the epithelium of the OV/IEO with a freehand line tool with line thickness set to match the thickness of the epithelium. Outlines were always drawn counterclockwise and were initiated at a point closest to the centroid of the aggregate. Intensity profiles were normalized to polar coordinates and intensity values averaged in 10° bins.

### Manipulation of RA signaling

Whole aggregates were exposed to various inhibitors of RA signaling and/or exogenous atRA at various points in the organoid culture protocol. RA activity was reduced by the pan-RAR inverse agonist AGN193109 (0.1 μM) or inhibitors to retinoic acid biosynthesis with WIN18446 (5 μM; Aldh1a2 specific), 673A (50 μM; Aldh1a family specific), or 646 (1 µM; Aldha1-3 selective). Inhibitors and atRA were applied in media exchanges at indicated time points between D7 and D16, during the phase of otic placode and otic vesicle induction and washed out when indicated by daily half-media exchanges. Vehicle for atRA, AGN193109, WIN18446, 673A, and 646 was dimethylsulfoxide at a final concentration of 0.1% or less

### Imaging

Brightfield imaging of aggregates and wholemounts, with or without chromogenic stains, was accomplished standard stereomicroscopes or Leica DM IL or DM LB compound microscopes outfitted with digital cameras (QImaging QICam or MicroPublisher0. Epifluorescence imaging was achieved with an Olympus BX51WI microscope and Orca-Flash 4.0 cooled-CCD camera (Hammamatus). Confocal imaging was performed using an Olympus FluoView 1000 or Leica SP8 Lightning microscope.

### Statistical analysis

Comparisons between 2 group means were performed using unpaired t-tests in Microsoft Excel. Comparisons amongst more than 2 groups were performed with one-way or two-way ANOVA and post-hoc tests in SPSS 24 or GraphPad Prism 10.

