## Supplemental Figures and Tables for "Retinoic acid signaling guides the efficiency of inner ear organoid-genesis and governs sensory-nonsensory fate specification"

**Document S1: Supplemental Information**

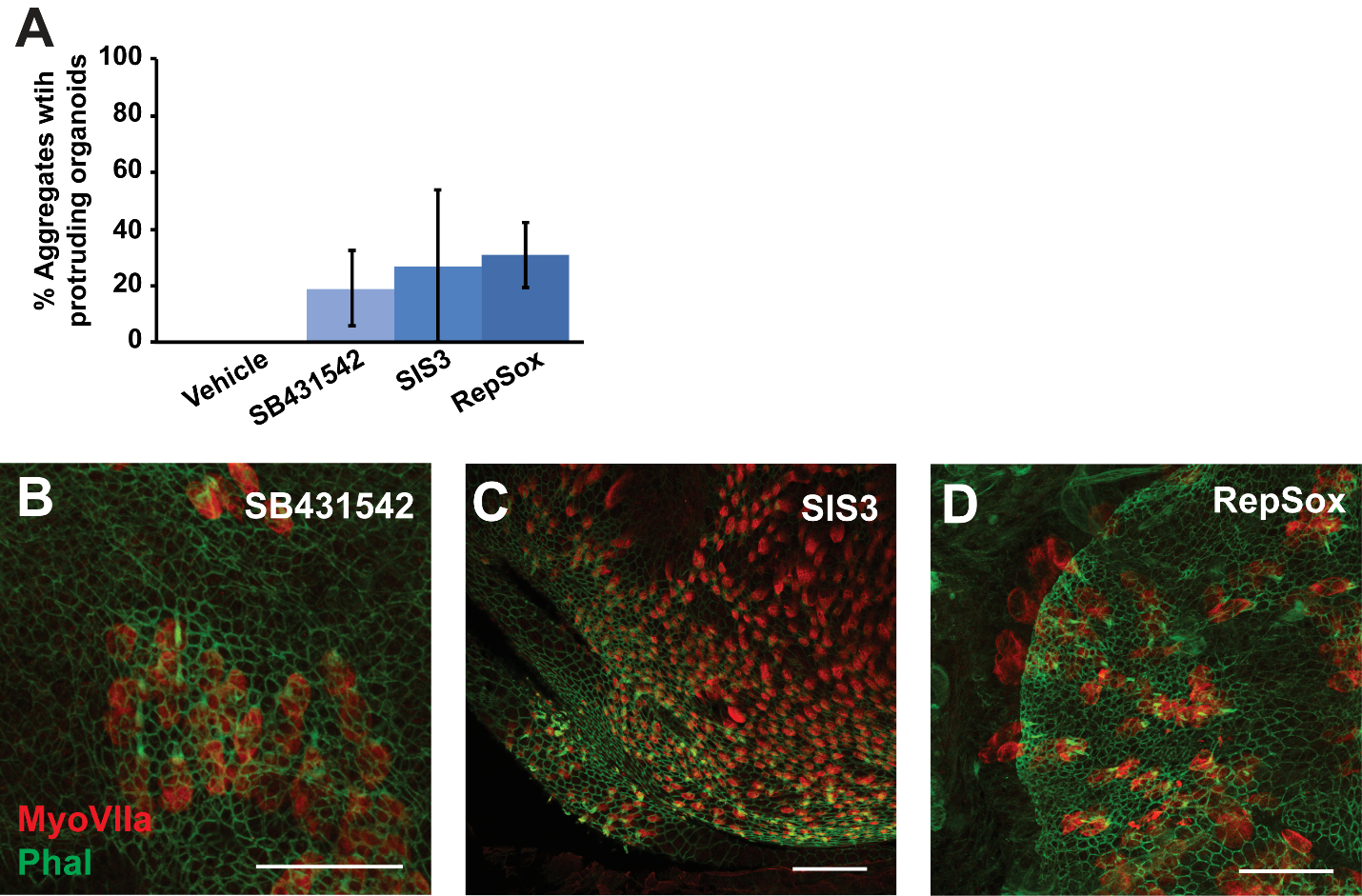

**Figure S1. Comparison of TGFβ pathway inhibitor effects on organoid production.** (A) The efficiency in production of hair-cell containing cystic organoids was quantified at D20 (*P*>0.05; mean ± one standard deviation). N = 8 (vehicle), 12 (SB431542), 7 (SIS3), 7 (RepSox). The vehicle control group never produced fluid-filled cysts (0/8 trials). (B-D) Example images of microdissected organoids from D20+ aggregates in each condition immunostained for MyoVIIa (red) and labeled with fluorescently tagged phalloidin (green) reveal hair cell production with all TGFβ inhibitors (0/3 vehicle control preparations showed MyoVIIa-positive hair cells). Scale bars: 50 μm.

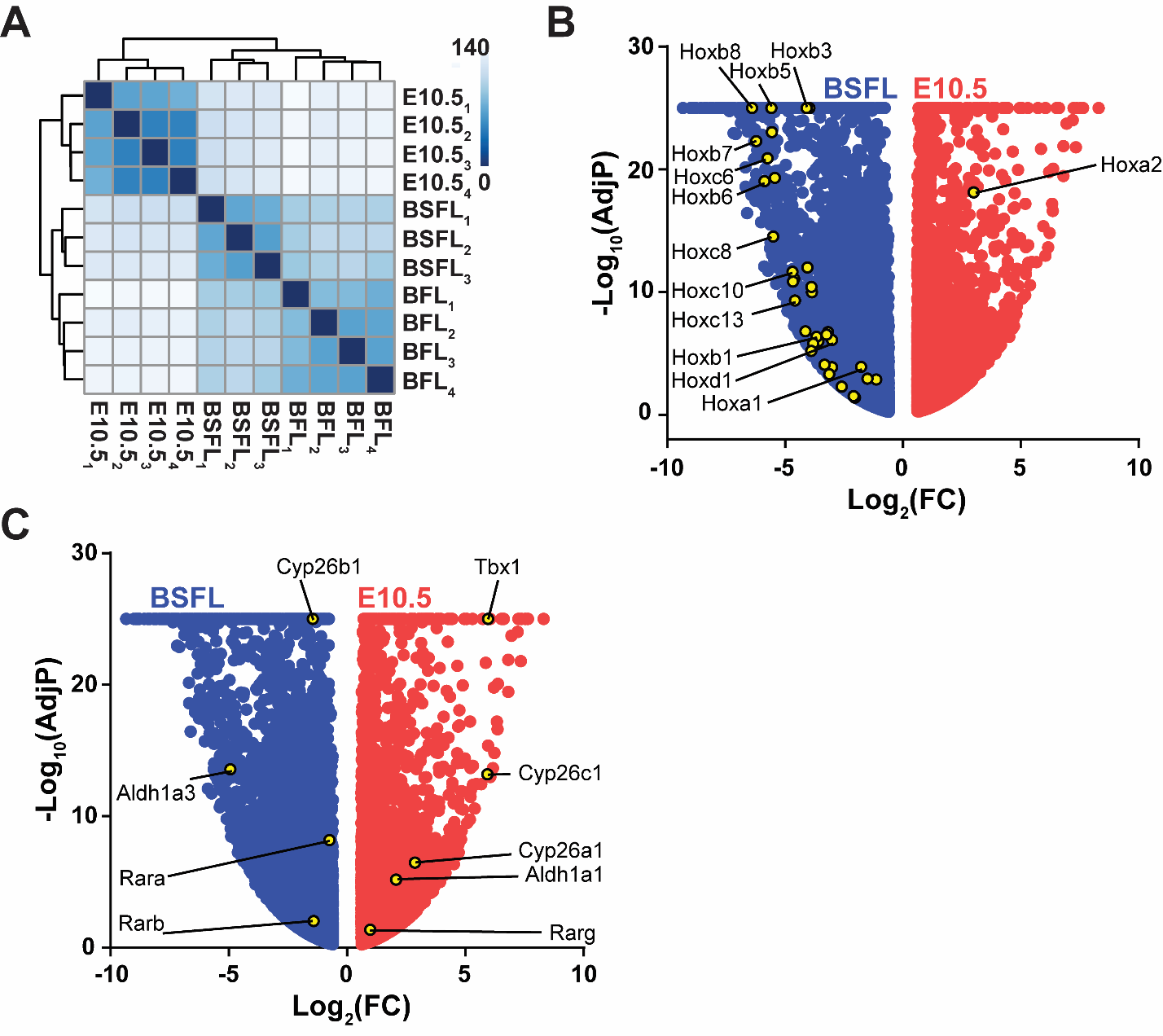

**Figure S2. RNASeq clustering and distribution of RA-related genes on volcano plots.** (A) Correlation heatmap and hierarchical clustering shows the greatest distance between *in vivo* and *in vitro* conditions, with secondary separation between the two culture conditions. Volcano plots are shown for E10.5 and BSFL DEGs with identification of (B) Hox genes and (C) other RA-related genes.

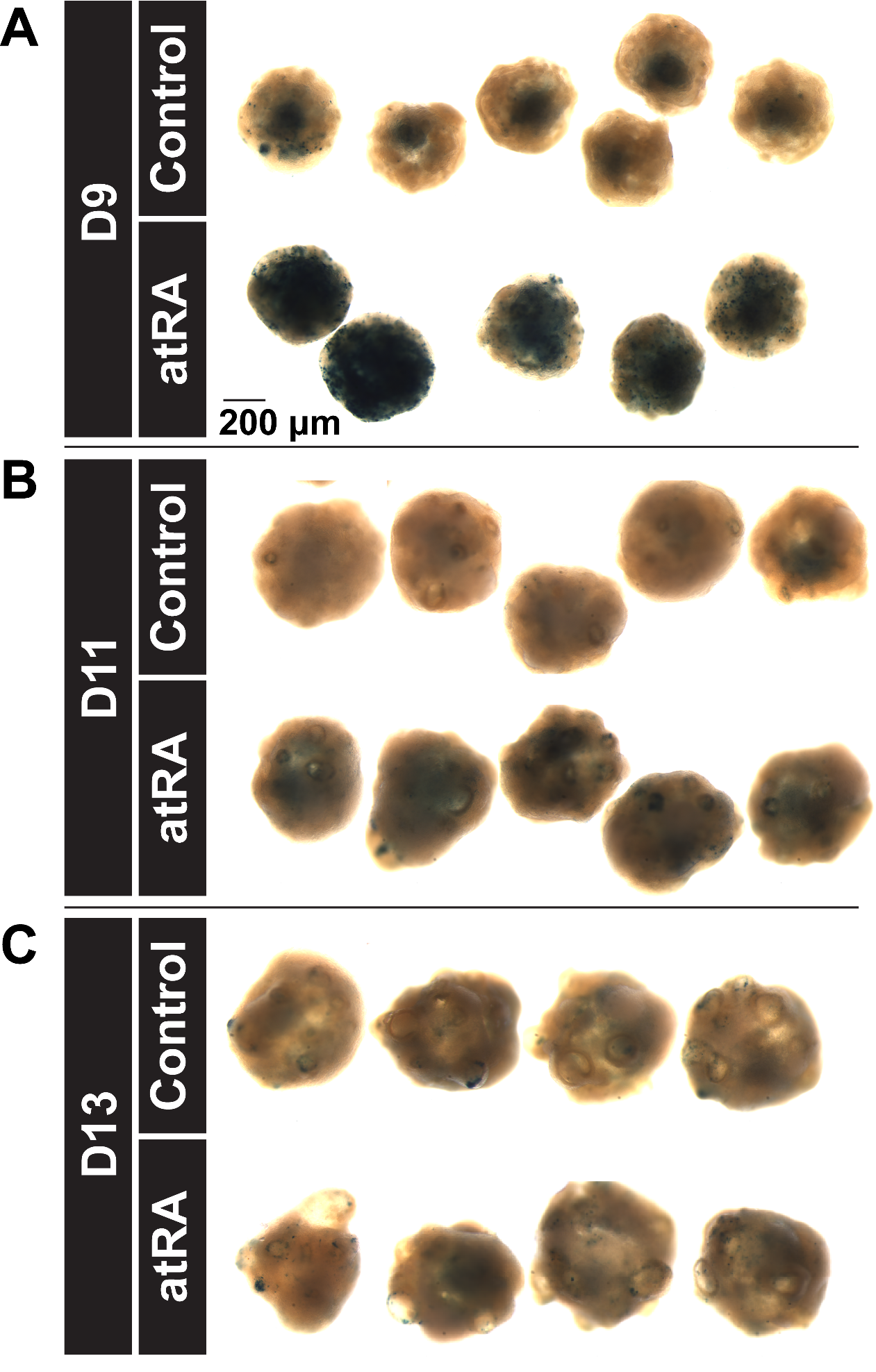

**Figure S3. Exposure to atRA increases lacZ reporter expression throughout RARE-lacZ aggregates at early time points.** Whole RARE-lacZ aggregates were stained with X-gal on (A) D9, (B) D11, and (C) D13 under control conditions and after application of 500 nM atRA 24-hours prior to fixation (i.e., D8, D10, or D12). Scale: 200 μm

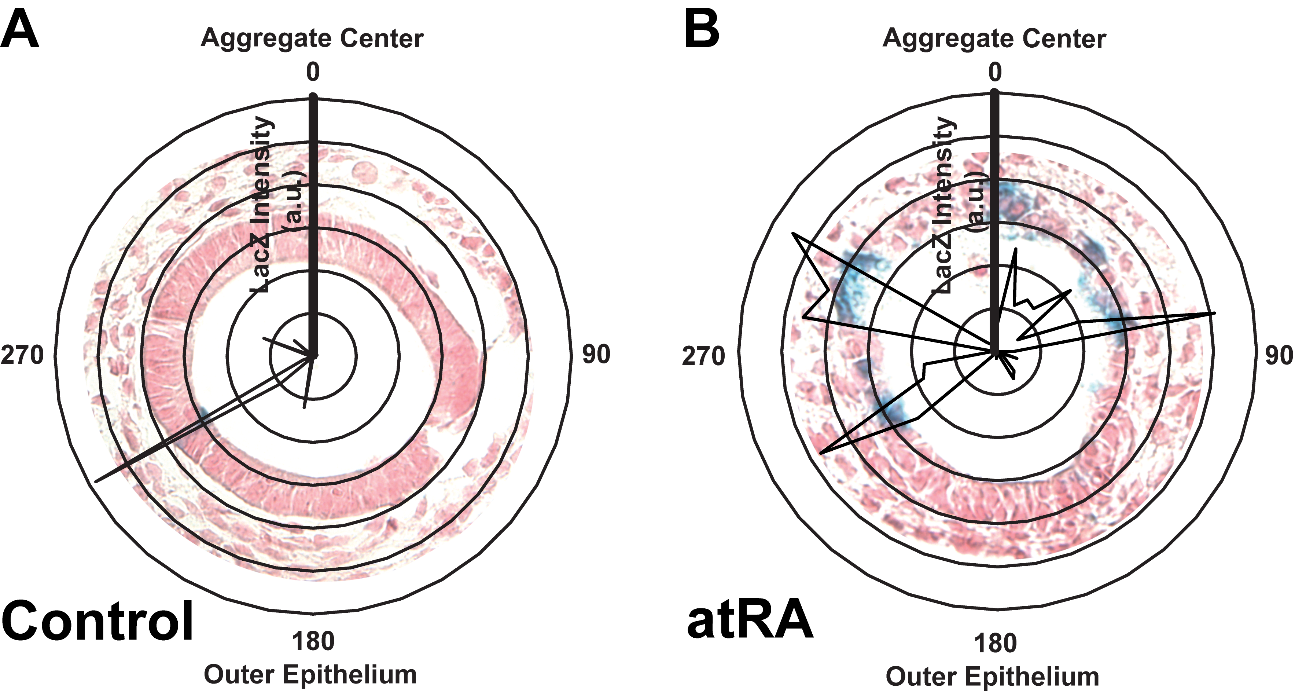

**Figure S4. Example quantification of Xgal-stained D13-14 otic vesicles.** Cryosections of D13-D14 aggregates stained with X-gal and counterstained with FastRed were imaged and analyzed for X-gal intensity as illustrated for (A) control and (B) atRA-treated cultures, where 500 nM atRA was applied from D8 to D12. Line intensity plots, in arbitrary units (a.u.) were translated to polar coordinates with the 0° oriented to the volumetric center of the aggregate.

**Table S1. Quality of RNA input and base call output**

| **Sample** | **Lane*** | **Total RNA quality (RIN)** | **PF Clusters (reads)** | **% PF Clusters** | **Yield (Mbases)** | **% >= Q30 bases** | **Mean Quality Score (Phred)** |
| --- | --- | --- | --- | --- | --- | --- | --- |
| BFL-1 | 1 | 8.5 | 24,280,297 | 91.05 | 1,578 | 93.33 | 36.2 |
| BFL-1 | 2 | 8.5 | 24,292,939 | 90.96 | 1,579 | 93.34 | 36.2 |
| BFL-2 | 1 | 8 | 21,681,310 | 91.01 | 1,409 | 93.03 | 36.14 |
| BFL-2 | 2 | 8 | 21,707,042 | 90.93 | 1,411 | 93.04 | 36.14 |
| BFL-3 | 1 | 8.4 | 24,874,381 | 91.12 | 1,617 | 93.05 | 36.14 |
| BFL-3 | 2 | 8.4 | 24,838,064 | 91.02 | 1,614 | 93.04 | 36.14 |
| BFL-4 | 1 | 8.2 | 27,530,756 | 91.19 | 1,789 | 93.22 | 36.18 |
| BFL-4 | 2 | 8.2 | 27,545,248 | 91.11 | 1,790 | 93.22 | 36.18 |
| BSFL-1 | 1 | 8.6 | 27,614,354 | 91.14 | 1,795 | 93.28 | 36.19 |
| BSFL-1 | 2 | 8.6 | 27,666,177 | 91.07 | 1,798 | 93.28 | 36.19 |
| BSFL-2 | 1 | 8.8 | 35,022,277 | 90.87 | 2,276 | 93.29 | 36.19 |
| BSFL-2 | 2 | 8.8 | 35,063,235 | 90.78 | 2,279 | 93.29 | 36.19 |
| BSFL-3 | 1 | 8 | 32,497,568 | 91.14 | 2,112 | 93.12 | 36.16 |
| BSFL-3 | 2 | 8 | 32,490,459 | 91.05 | 2,112 | 93.11 | 36.16 |
| E10-1 | 1 | 9.7 | 30,318,211 | 91.11 | 1,971 | 93.22 | 36.18 |
| E10-1 | 2 | 9.7 | 30,332,421 | 91.03 | 1,972 | 93.21 | 36.18 |
| E10-2 | 1 | 9.8 | 27,370,770 | 91.02 | 1,779 | 93.16 | 36.17 |
| E10-2 | 2 | 9.8 | 27,377,820 | 90.94 | 1,780 | 93.16 | 36.17 |
| E10-3 | 1 | 10 | 29,853,525 | 90.35 | 1,940 | 93.21 | 36.18 |
| E10-3 | 2 | 10 | 29,877,200 | 90.26 | 1,942 | 93.21 | 36.18 |
| E10-4 | 1 | 9.1 | 28,635,237 | 90.63 | 1,861 | 93.23 | 36.18 |
| E10-4 | 2 | 9.1 | 28,648,774 | 90.51 | 1,862 | 93.22 | 36.18 |

* Each sample was run in 2 lanes

**Table S2. Alignment of BAM file reads to annotated transcripts from HTSeq Count**

| **Sample** | **Total BAM file reads (millions)** | **% Mapped** | **% Mapped to forward strand** | **Reads mapped but no feature annotated (millions)** | **Ambiguous Reads with >1 feature at the mapped locus (millions))** | **Reads not mapped to a locus (millions)** | **Unique reads**  **Total BAM file reads - No feature - Ambiguous - Not aligned (millions)** | **% Unique reads** | **Alignment not unique**  **Total alignments of reads mapped to >1 locus (millions)** |
| --- | --- | --- | --- | --- | --- | --- | --- | --- | --- |
| BFL-1 | 79.1 | 97.7 | 51.4 | 3.7 | 1.5 | 1.8 | 72.2 | 91.2 | 43.1 |
| BFL-2 | 54.6 | 96.9 | 50.3 | 3.0 | 1.8 | 1.7 | 48.2 | 88.2 | 18.2 |
| BFL-3 | 80.3 | 97.6 | 46.8 | 3.2 | 1.6 | 2.0 | 73.5 | 91.6 | 43.4 |
| BFL-4 | 69.2 | 96.7 | 49.7 | 4.5 | 2.2 | 2.3 | 60.2 | 87.1 | 23.0 |
| BSFL-1 | 68.8 | 96.5 | 47.5 | 4.5 | 2.2 | 2.5 | 59.6 | 86.6 | 22.4 |
| BSFL-2 | 110.8 | 97.5 | 52.9 | 4.6 | 2.2 | 2.8 | 101.2 | 91.3 | 58.5 |
| BSFL-3 | 81.1 | 96.6 | 49.5 | 4.7 | 2.4 | 2.8 | 71.2 | 87.9 | 26.6 |
| E10-1 | 74.0 | 97.3 | 61.1 | 3.3 | 2.4 | 2.0 | 66.3 | 89.6 | 21.9 |
| E10-2 | 67.4 | 97.3 | 46.6 | 3.1 | 2.2 | 1.9 | 60.2 | 89.4 | 20.6 |
| E10-3 | 73.8 | 97.3 | 49.9 | 3.1 | 2.5 | 2.0 | 66.2 | 89.7 | 22.9 |
| E10-4 | 70.1 | 97.3 | 53.4 | 3.2 | 2.2 | 1.9 | 62.8 | 89.5 | 21.1 |
